# Optimization of PURE system composition using automation and active learning

**DOI:** 10.64898/2026.03.23.713685

**Authors:** Yannick Bernard-Lapeyre, Céline Cleij, Andrei Sakai, Marie-José Huguet, Christophe Danelon

## Abstract

Protein synthesis using recombinant elements (PURE) system has been widely applied in various biological research fields and synthetic cell construction. Optimization efforts to enhance the PURE system performance by adjusting its individual components have remained limited to the expression of single genes with a small number of molecular compositions tested, making it difficult to link component composition to system-level performance across different DNA contexts. Here, we combine automated acoustic liquid handling with an active learning framework to explore broadly the compositional landscape of PURE system. By grouping the 69 individual components (including proteins and tRNAs) into 21 functional sets and iteratively guiding experiments with active learning, we rapidly identify improved compositions and demonstrated up to 3-fold enhancement in protein yield and translation rate for a single reporter gene. We further show that optimization drivers differ between low and high DNA concentrations, revealing that optimal PURE compositions are DNA concentration-dependent. We then apply this optimization strategy to enhance the expression of a 41-kb synthetic chromosome containing 15 genes by maximizing the fluorescence intensities of two reporter proteins. While a 3-fold improvement could be reached on the two gene products guiding learning, a full proteomic analysis revealed that optimization is gene-specific, i.e., changes in PURE system compositions differently impact the amounts of synthesized proteins encoded on the same DNA template. Together, this work establishes active learning as an efficient strategy to navigate the high-dimensional PURE compositional space and provides mechanistic insight into DNA context-dependence of gene expression optimization.

## INTRODUCTION

Cell-free expression (CFE) systems have become versatile tools for synthetic biology, enabling rapid testing of genetic circuits, metabolic pathway engineering, and bottom-up synthetic cell construction^1–3^. Among these systems, the protein synthesis using recombinant elements (PURE) system stands out for its strictly defined composition and absence of nucleases and proteases which are typically present in cell lysates^4–6^. By reconstituting transcription, translation, energy regeneration, and tRNA charging from purified components, PURE system provides a minimal and controllable environment for gene expression. Essential cellular functions driven by protein synthesis with PURE system have been reconstructed within liposomes, including DNA replication, lipid synthesis and cell division^7–9^. PURE system has also been leveraged for applications such as cell-free screening of antimicrobial peptides, allowing for rapid and controlled assessment of peptide activity *in vitro*^10^. Other fields of application include protein solubility and chaperone effects^11,12^, or the synthesis of complex membrane proteins^13^, incorporation of unnatural amino acids^14^, and protein engineering through directed evolution^15^.

However, CFE with current PURE system composition is constrained by a modest protein yield, slow translation rate, and short reaction lifetime, particularly when compared to *E. coli*-based extract systems. In previous efforts to boost PURE performance, substrate feeding strategies have been explored and key translation factors have been identified as limiting factors^16,17^. Moreover, alternative energy regeneration mechanisms have been implemented^18^ and improved tRNA recycling has been attempted^19^. While these studies revealed that individual factors can influence performance under certain conditions, they also highlighted the strong context dependence of these improvements and the difficulty of predicting beneficial modifications *a priori*^17–19^. Therefore, rational optimization of PURE reaction efficiency remains a challenge, and better combinations of biochemical components can be hard to find without running many experiments.

Similar shortcomings exist with lysate-based CFE systems. To overcome them, recent studies have leveraged design of experiment approaches, machine learning and laboratory automation to efficiently explore the high-dimensional parameter space of CFE systems with a focus on reaction buffer optimization^20–23^. Using an automated workflow combining Echo acoustic liquid handling and an active-learning framework implemented in a Google Colab notebook called METIS^22^, Pandi et al. optimized the buffer composition of an *E. coli*-based lysate for improving protein production yield, genetic circuit performance (LacI-based multi-level controllers) and for metabolic pathway optimization (CETCH cycle). METIS couples XGBoost regression^24^ with Bayesian optimization^25^ to guide the systematic exploration of multicomponent biological networks. This method has recently been applied to enhance expression in various lysate-based systems^26,27^. However, in these studies, only the buffer composition was modified, while the macromolecular content was a fixed ’black box’, thereby limiting the exploration space and missing opportunities for further enhancement. A notable exception is Contreras-Llano et al. (2020)^28^, who increased the protein content of their CFE system by overexpressing 34 core translation factors in their source bacteria, resulting in up to a five-fold increase in cell-free protein synthesis capacity. Such findings, combined with the known epistatic interactions among the purified elements of the translation machinery^29^, suggest that further optimization would be possible by scanning the composition of the protein hardware itself.

Here, we perform a PURE system-wide optimization, tuning not only substrates and cofactors but also major macromolecular machineries such as aminoacyl-tRNA synthetases, tRNAs, translation factors, and ribosomes. Using iterative active learning and automated experimentation, we first demonstrate that optimized PURE compositions can yield up to three-fold improvement of synthesized proteins for a single-gene reporter. The components that most impact the yield differ substantially between low and high concentrations of the input DNA, revealing distinct mechanistic regimes. Finally, we generalize this workflow to enhance the expression of a 41-kb synthetic chromosome encoding 15 genes. We find that PURE optimization is strongly context-dependent, with DNA concentration–dependent constraints and gene-specific trade-offs that limit chromosome-scale expression enhancement. These results establish an extensible framework for improving PURE system and unveil unforeseen challenges toward user-defined chromosome-scale gene expression profiles *in vitro*.

## RESULTS

### Defining the parameter space for PURE system optimization

To facilitate exploration and reduce the number of variables in the active learning process, while also minimizing component preparation time, we grouped the 69 individual components present in PURE system into 21 functional groups. Six components remained as individual variables: HEPES-KOH (hepes), potassium glutamate (k-glut), glutathione (gsh), spermidine, 10-formyl-tetrahydrofolate (fthf), creatine phosphate (cp), tyrosine (tyr), cysteine (cys), T7 RNA Polymerase (t7pol), and ribosomes (**Table 1**). tRNAs are already provided as a mix from the supplier. The remaining components were clustered into 15 sets based on their biochemical functions. For instance, all initiation factors were pooled together, as well as all aminoacyl-tRNA synthetases.

**Table 1:**
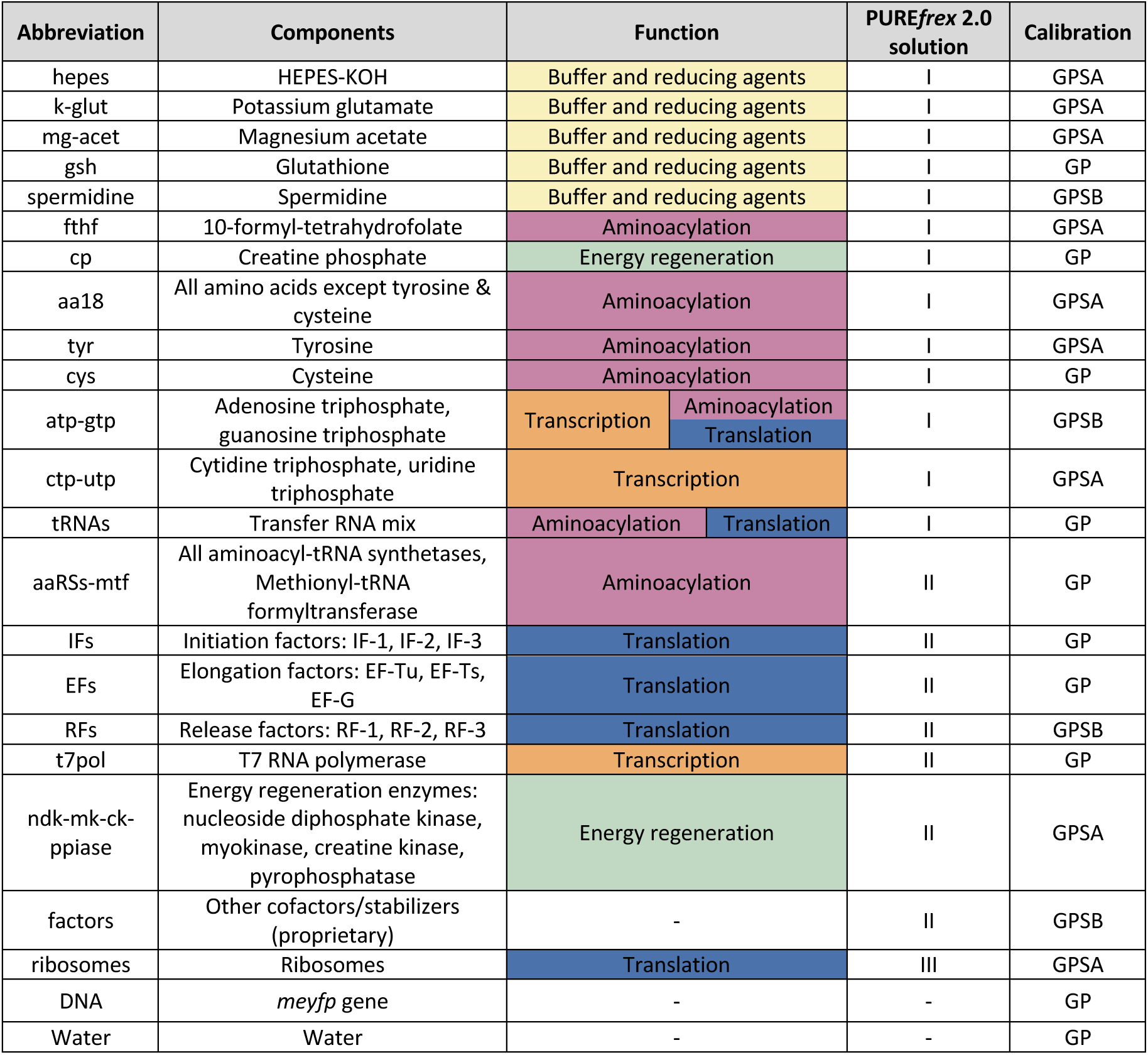
List of PURE system components with abbreviations, functional classification and Echo calibration type used for dispensing. GP category is for low-high viscosity fluids (<50% glycerol). GPSA category is for low-high viscosity fluids (<50% glycerol) with low level surfactants. GPSB category is for low-high viscosity fluids (50% glycerol) with high level surfactants.

We performed calibration tests with the Echo liquid dispenser for automated distribution of PURE components onto microplates (**Figure 1**). This step was crucial, as the PURE components contain varying amounts of glycerol, a factor that influences dispensing accuracy. We performed a combined analysis of droplet size/morphology on a filmed plate and volume measurements in the source wells. Based on this assessment, we selected the most appropriate calibration type from the five available options with the source plate used, for each of the 21 sets of components (**Table 1**). This also allowed us to determine how much extra volume was to be added for each component to account for dispensing errors, which is highly dependent on solution viscosity. Besides calibration, we optimized handling protocols for these components, considering temperature, dilution and preparation of working solutions of the components (Methods). Solution I components (feeding mix) were handled at room temperature and were the first to be distributed with Echo when assembling reactions due to the tendency of some components to precipitate when handling under cold conditions. Order of the assembly proved to be important with initial assembly attempts resulting in precipitation or hampered activity. Proteins (Solution II) were prepared on ice to preserve activity. Following Echo’s requirements of the source plate being at room temperature, the plate was not filled on ice but at room temperature. Despite the components not being cooled before and during Echo distribution, PURE reactions still showed activity. Leftover components were flash-frozen immediately after dilution and remained functional for later experiments.

**Figure 1:**
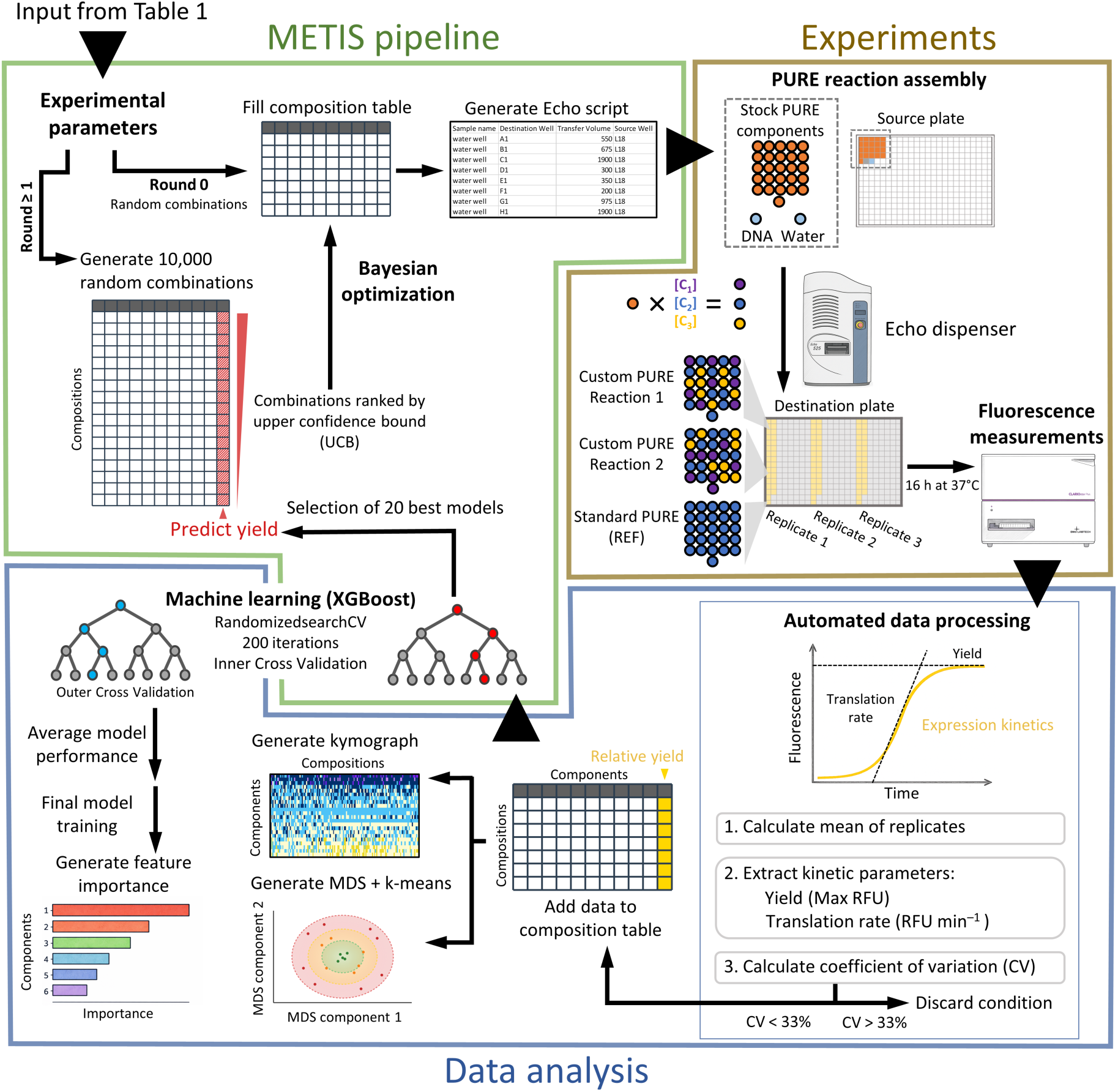
Active learning workflow for PURE system optimization. The 69 PURE components are grouped into 21 sets, and experimental parameters and concentration ranges are defined in METIS. METIS generates initial compositions (Round 0) and converts dispensing instructions into an Echo-compatible .csv script. Components (including two reporter genes) with calibrated liquid classes are loaded onto a 384-well source plate and reactions are assembled with the Echo instrument. Real-time fluorescence kinetics are measured and fitting with a phenomenological model is used to extract the maximum yield and apparent translation rate. From Round 1 onward, METIS trains an ensemble of XGBoost regressors on accumulated data, using the 20 best models to predict yields for 10,000 candidate compositions. Bayesian optimization ranks candidates using an Upper Confidence Bound (UCB) metric, and the top conditions are selected for the next round.

To scan PURE compositions, we chose three concentrations for each group of components (0.5×, 1×, 2× their concentration in regular PURE*frex*2.0), corresponding to a combinatorial space of 3^21^, i.e., over 10 billion possible mixtures. As the entire compositional space is too large for systematic exploration, we opted for an active learning-assisted optimization approach to accelerate the search. To measure performance, we performed a dual reporter assay, combining the expression of 100 pM *meyfp* and 10 pM *lacZ* genes, both in the form of linear DNA templates (PCR products) in single reactions. However, active learning was solely based on mEYFP fluorescence kinetics. Expression of *lacZ* was quantified as an endpoint measurement of β-galactosidase activity using a change of absorbance after addition of chlorophenol red-β-D-galactopyranoside (CPRG) substrate and its conversion into chlorophenol red (CPR). The inclusion of a second reporter allowed us to evaluate whether changes in PURE composition similarly influenced the co-expression of another gene.

### Validation of Echo-based PURE assembly

To validate the use of acoustic liquid handling for PURE system assembly, we first calibrated dispensing parameters for all 21 component groups to account for differences in solution viscosity and glycerol content, and established handling protocols that preserved component stability and activity (**Table 1**). Using these conditions, we reconstituted the reference PURE*frex* 2.0 composition by Echo dispensing and compared its performance to manually assembled reactions and to the commercial kit (Solutions I, II and III). Echo-assembled and manually assembled PURE reactions displayed highly similar *meyfp* expression kinetics, yields (maximum fluorescence), and apparent translation rates (maximum slope of a fitted expression curve), demonstrating that automated dispensing does not impair PURE activity and that the system can be robustly reconstituted from individually prepared components (**Figure S1**). The commercial PURE*frex* 2.0 kit consistently produced slightly higher yields, indicating that formulation, storage, and handling of PURE constituents influence absolute performance. Comparison across two independently prepared batches of PURE components further revealed substantial batch-to-batch variability in overall activity, while preserving relative agreement between Echo- and manually-assembled reactions at two different DNA concentrations (**Figure S1**). Together, these results validate Echo-based automation as a reproducible method for assembling PURE reactions, while indicating batch-dependent effects.

### Active learning improves expression yield

We then sought to discover new compositions of PURE system with enhanced expression yield of *meyfp* (0.1 nM) compared to the standard composition, thereafter called reference (REF), assembled with the Echo dispenser. In Round 0, we established a first dataset using 28 random combinations generated by METIS, alongside the REF composition, all tested in triplicate (**Figure 1** and **Figure 2A,B**). Three conditions were discarded due to a high coefficient of variation (CV>33%) between the technical replicates. Among the 25 remaining compositions, three outperformed the REF, with the best-performing condition reaching a relative yield of 1.7 (**Figure 2C**). These data were fed back into the algorithm, using the value of the relative yield as the parameter to be maximized. New conditions were generated and tested for round 1, and the data of both rounds 0 and 1 were used to generate conditions for round 2 (**Figure 2A**). The two highest performing conditions had relative yields of 2.4 and 2.1 (**Figure 2C**). From rounds 0 to 2, iterative optimization through METIS led to a marked improvement in fluorescence levels, as seen with the higher incidence of kinetic profiles outperforming REF (green curves, **Figure 2B**). This result underpins that even though the range of concentration changes is narrow (0.5 to 2), the effects on yield can be substantial even after only two rounds of optimization.

**Figure 2:**
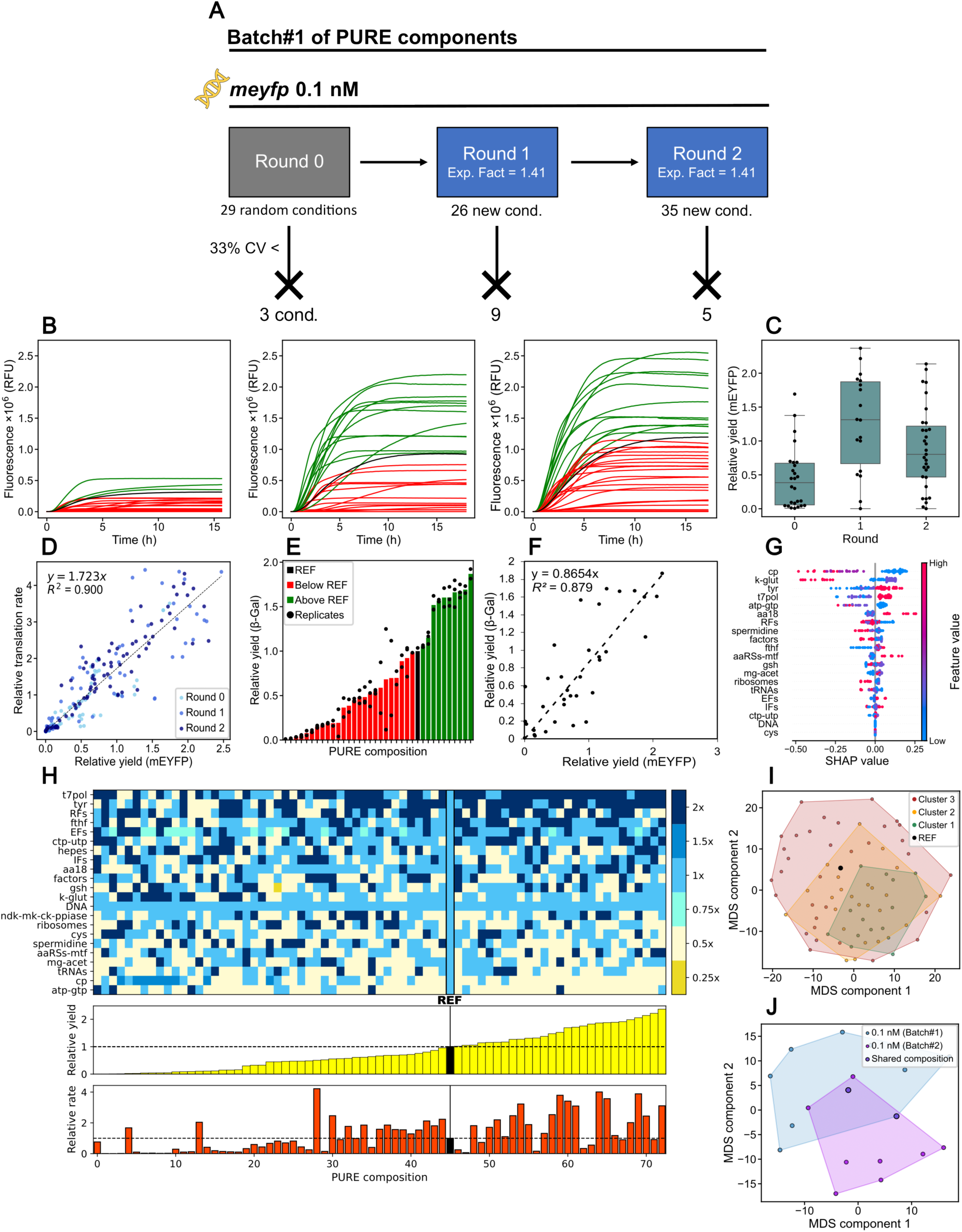
Active learning improves yields across rounds. **(A)** Overview of conditions tested in rounds 0–2. Conditions with CV > 33% were discarded. The exploration factor is indicated for each active learning round. **(B)** Mean mEYFP fluorescence kinetics for all conditions. DNA concentration was 100 pM for *meyfp* and 10 pM for *lacZ*. Each curve is the mean of n = 2 or 3 technical replicates. Black, REF composition; green, curves with final yield > REF; red, curves with final yield < REF. **(C)** Distribution of relative yields (max fluorescence normalized to REF) across the three rounds. Box plots show median (center line), interquartile range (IQR, box), and 1.5× IQR (whiskers), with individual data points overlaid. **(D)** Correlation between relative translation rate and relative yield across rounds. Data points represent individual replicates. The linear fit and corresponding R² are shown. **(E)** Relative β-galactosidase yields derived from CPRG conversion assays. Mean of 2 replicates in green: >REF; red: <REF. Individual replicates represented by black points **(F)** Correlation between mEYFP and β-galactosidase relative yields (round 2). The linear fit and corresponding R² are shown. **(G)** Shapley values (SHAP) beeswarm plot showing components most influencing yield prediction. Each point corresponds to one experimental condition. The x-axis shows the SHAP value (positive values increase, negative values decrease the predicted yield); colors indicate relative component concentrations within the range tested (blue, low; red, high). **(H)** Kymograph of all tested compositions sorted by relative yield mEYFP (subplot, top). The corresponding relative translation rate values are displayed in the subplot beneath. Color scale depicts component concentrations relative to the REF composition. Dashed lines in the subplots indicate values of 1 (REF). **(I)** Multidimensional scaling (MDS) representation (Manhattan distances) of all conditions with k-means clustering (k = 3). Colors represent clusters ranked by relative mEYFP yield, with Cluster 1 in green containing top performing conditions. **(J)** MDS comparison of top 10 conditions from two PURE component batches (Batch#1 and Batch#2), showing partial overlap but batch-specific signatures.

We performed a multidimensional analysis of PURE kinetics across the three rounds of active learning by plotting the relative apparent translation rate of mEYFP against the relative yields. A strong positive correlation was observed, indicating that compositions leading to higher translation rates also produce more proteins (**Figure 2D**). This trend remains consistent across all three experimental rounds. Overall, the finding that translation rate and end-point protein yield are positively correlated suggests that optimization based on one kinetic parameter (maximizing yield) may lead to concurrent improvement of the other (maximizing rate).

The expression level of the second reporter gene, *lacZ*, was assessed by measuring the initial change of CPR absorbance upon addition of CPRG in pre-ran PURE reactions containing steady amounts of the synthesized β-galactosidase. The change in absorbance was measured for each composition, and the steepest slope was extracted and normalized to that of the REF composition, giving relative yields of β-galactosidase as a performance metric (the CPRG conversion rate is proportional to the concentration of synthesized enzyme) (**Figure 2E and S2)**. Plotting the two metrics of PURE activity (relative yield of mEYFP and relative yield of β-galactosidase) for all Round 2 conditions reveals a positive correlation (**Figure 2F**), suggesting that optimization of PURE composition (learning through mEYFP fluorescence intensity) may apply to co-expressed genes as well.

To evaluate learning accuracy and the intrinsic predictability of the dataset across all rounds, we trained XGBoost regression models using a nested cross-validation framework. An outer cross-validation loop was used to estimate generalization performance, while an inner loop optimized model hyperparameters. Predicted relative yields from the held-out outer test folds were compared to experimentally measured values, and performance metrics were aggregated across folds (**Figure S3A**). The model captured limited but non-random structure in the data (R² = 0.16 ± 0.24), consistent with the small size of the dataset. This indicates that the model seizes coarse yield trends and relative ranking rather than accurate values.

The trained models were subsequently used for feature importance analyses to identify PURE components whose concentration changes most strongly influenced the yield. Three complementary feature importance metrics were employed: XGBoost gain importance, SHAP (Shapley additive explanations) values, and permutation importance (**Figure S3B**). A consensus importance score was calculated by normalizing each metric and averaging them. Because this consensus closely resembled the SHAP profile, SHAP values were selected as the primary metric for interpretation (**Figure 2G**). SHAP analysis identified cp and k-glut as the most influential components when decreased. Tyr and t7pol also showed influence on prediction when increased.

To visualize compositional trends, we plotted all tested compositions from rounds 0–2 as a kymograph, sorted by relative yield and ordered by total component concentration from high (top) to low (bottom). Relative yield of each condition as well as the relative translation rates were also added as subplots. This analysis revealed enrichment of t7pol and tyr in high-yielding compositions relative to REF, whereas atp-gtp, cp, and tRNAs were generally depleted (**Figure 2H**).

Finally, similarities among PURE compositions were examined using multidimensional scaling (MDS) combined with k-means clustering (k = 3). Conditions were projected onto a two-dimensional space in which distances reflect compositional dissimilarity. K-means clustering with convex hulls delineated three distinct groups associated with relative yield. The highest-performing compositions (Cluster 1; 1.47 ≤ relative yield ≤ 2.37) formed a compact cluster, indicating high compositional similarity. In contrast, the two lower-performing clusters (Cluster 2; 0.61 ≤ relative yield ≤ 1.38; Cluster 3; 0 ≤ relative yield ≤ 0.57) exhibited substantially greater heterogeneity (**Figure 2I**).

We validated the workflow on a separately purified batch of PURE components (Batch#2) by testing 110 new combinations and 20 “most informative” conditions identified by METIS from earlier rounds (**Figure S4A**). Absolute fluorescence of the REF condition was drastically reduced in this batch, leading to inflated relative yields (**Figure S4B**). Two compositions achieved relative mEYFP yields of 27 and 21 within three optimization rounds, and a strong positive correlation between yield and apparent translation rate was confirmed (**Figure S4C,D**). The method proved reproducible across experimental days and DNA template batches, with consistent outcomes for 14 tested conditions (**Figure S4E**).

Using the same nested cross-validation framework, XGBoost models trained on Batch#2 exhibited very low predictive power (R² = 0.01 ± 0.26) (**Figure S5A**), i.e., differences in yield are largely not predictable from the component concentrations alone. Despite this poor predictive power, feature importance analysis remains informative: it indicates which components most influence the model’s decisions, revealing compositional trends and potential drivers of yield even when exact predictions are unreliable. Here, it consistently identified t7pol as the determining variable to model decisions, followed by tRNA, atp-gtp, and mg-acet, although their relative rankings varied across methods (**Figure S5B**). These results indicate that, for this batch, the data contain insufficient structured information for reliable yield prediction, and feature importance should be interpreted only qualitatively.

Kymograph analysis identified mainly t7pol and tyr as enriched in high-yielding conditions, while cp, atp-gtp, tRNAs, and ribosomes were strongly depleted (**Figure S4H**). Multidimensional scaling combined with k-means clustering (k = 3) again revealed distinct compositional groupings (**Figures 2I and S4G**). On the two-dimensional map, high-yielding conditions (Cluster 1; 9.01 ≤ relative yield ≤ 27.32) occupied a narrow space, indicating conserved optimal composition features within this batch. In contrast, the lower-performing clusters (Cluster 2; 3.10 ≤ relative yield ≤ 8.52; Cluster 3; 0 ≤ relative yield ≤ 3.03) displayed greater compositional heterogeneity (**Figure S4H**). The ten highest-yielding compositions from Batch#1 and Batch#2 were compared using MDS. Partially overlapping clusters emerged, indicating that while optimal formulations share conserved features across conditions, batch-to-batch variability remains a significant source of divergence (**Figure 2J**).

### Optimized PURE compositions differ at low and high DNA concentrations

Having obtained enhanced PURE compositions for expression of low-concentration (0.1 nM) DNA, we applied the active learning workflow to improve protein yield from 2 nM *meyfp* DNA, asking whether the factors limiting performance depend on DNA concentrations^19^. The experiments were performed using Batch#2. Over three rounds of active learning (denoted rounds 6-8, **Figure 3A**), together scanning 136 new and 20 ’most informative’ combinations from previous rounds at 0.1 nM DNA, new compositions were discovered with relative yields > 3, with the best-performing one reaching a value of 3.8 (**Figure 3B,C**). Scatter plot of relative translation rate versus relative yield of mEYFP (**Figure 3D**) reveals positive correlation, albeit the strength of the correlation diminishes at higher values, a trend that was less pronounced at 0.1 nM DNA (**Figure 2D**).

**Figure 3.**
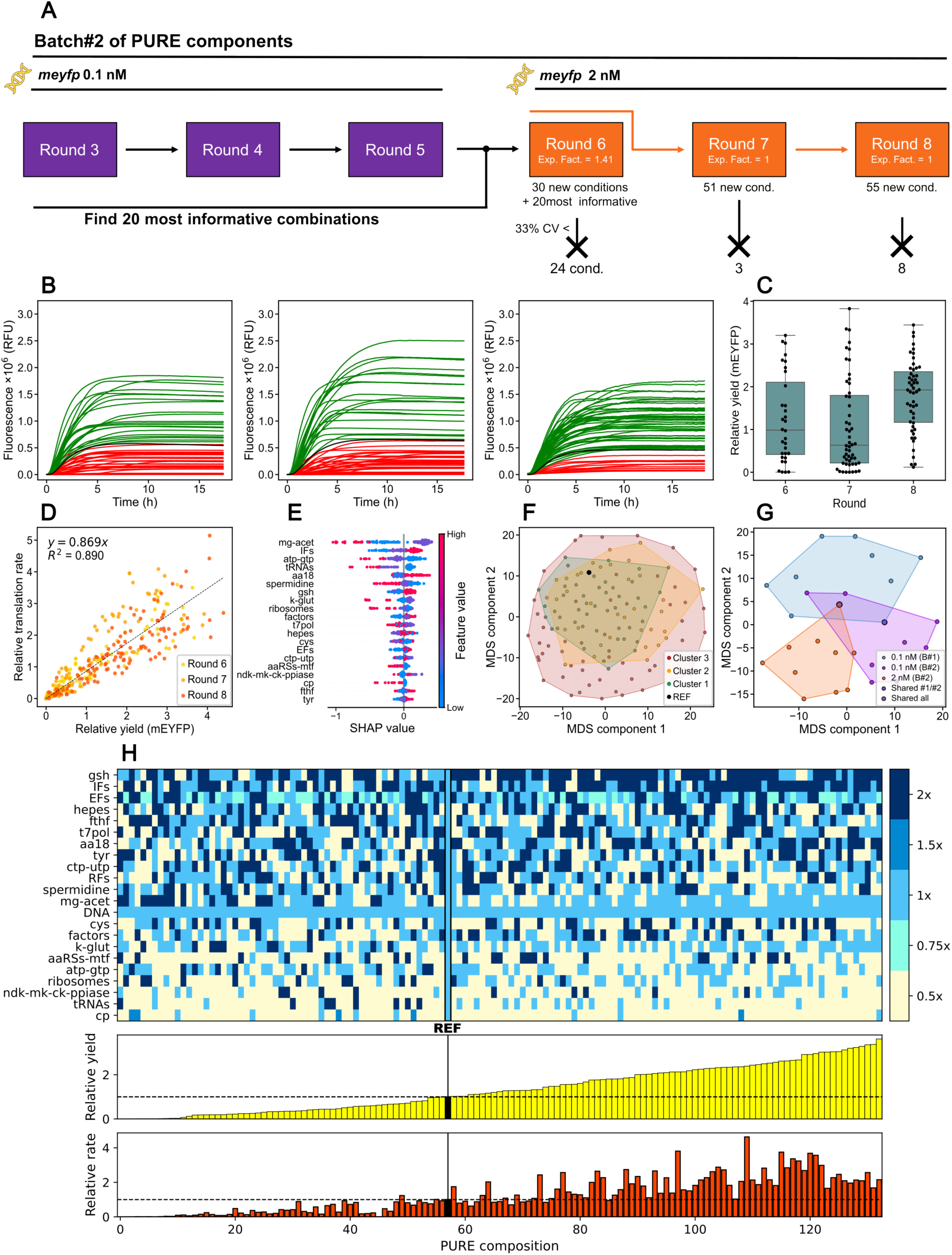
Distinct factors drive PURE optimization at low vs. high DNA concentration. **(A)** Overview of the PURE optimization strategy at 2 nM *meyfp* with indicated conditions generated and discarded when CV > 33% across technical replicates. Data from previous rounds were used to generate round 6, in addition to 20 most informative combinations from rounds 3 to 5. Rounds 7 and 8 were generated from data at 2 nM DNA only. The exploration factor is indicated for each round. **(B)** Mean mEYFP fluorescence kinetics for all conditions tested across rounds 3–5. Each curve is the mean of n = 2 replicates. Black, REF composition; green, curves with final yield > REF; red, curves with final yield < REF. **(C)** Distribution of relative yields for rounds 6–8. Box plots show median (center line), interquartile range (IQR, box), and 1.5× IQR (whiskers), with individual data points overlaid. **(D)** Correlation between relative translation rate and relative yield of mEYFP across rounds. The linear fit and corresponding R² are shown. **(E)** Shapley values (SHAP) beeswarm plot showing components most influencing yield prediction. **(F)** MDS representation (Manhattan distances) of all conditions with k-means clustering (k = 3). Colors represent clusters ranked by relative mEYFP yield, with cluster 1 in green containing top performing conditions. **(G)** MDS analysis of the ten highest-yielding compositions from all datasets (Batch#1 and Batch#2 at 0.1 nM DNA, and Batch#2 at 2 nM DNA). **(H)** Kymograph of all tested compositions from rounds 3–5 sorted by relative yield mEYFP (subplot, top). The corresponding relative translation rate values are displayed in the subplot beneath. Color scale depicts component concentrations relative to the REF composition. Dashed lines in the subplots indicate values of 1 (REF).

To assess predictive power at high DNA load, we applied the same nested cross-validation framework to the dataset obtained at 2 nM DNA. Predictive power improved substantially compared to 0.1 nM DNA, reaching a mean coefficient of determination R² = 0.34 ± 0.29 (**Figure S6A)**. Feature importance analyses were consistent across methods, converging on mg-acet as the main determinant to yield variation (1× concentration), followed by IFs, atp-gtp, tRNAs, and spermidine (**Figures 3E and S6B**). Together, these results indicate that, at high DNA concentration, PURE behavior becomes more predictable.

This result is corroborated by kymograph analysis revealing a pattern of component concentrations leading to a relative yield > 1 that is notably different than the one observed at lower DNA quantity (**Figure 3H**). For instance, compositions associated with a high relative yield contain higher concentrations of gsh, IFs and EFs, lower concentrations of cp, tRNAs and ndk-mk-ck-ppiase compared to REF, and a concentration of mg-acet strongly set at 1×. Interestingly, t7pol also remained at standard concentration, whereas it was doubled at 0.1 nM DNA.

MDS combined with k-means clustering (k = 3) identified three compositional clusters (**Figure 3F**). Cluster 1 (2.006 ≤ relative yield ≤ 3.615) and cluster 2 (0.9624 ≤ relative yield ≤ 1.873) occupied relatively compact regions of MDS space, whereas cluster 3 (0.002136 ≤ relative yield ≤ 0.8222) spanned a broader area.

The combined MDS map of all three experimental sets (two batches of reagents at 0.1 nM DNA and one batch at 2 nM DNA) revealed a small region of overlap among the clusters, potentially defining a compositional space for a generic, enhanced PURE system (**Figure 3G**). Notably, the MDS analysis indicated that the top-performing compositions at 0.1 nM and 2 nM occupy distinct regions, supporting the idea that the system operates under different constraints.

### Validation of optimized conditions through manual assembly

Next, we tested whether enhanced PURE compositions (ePURE) identified by the active learning-assisted Echo workflow retain their performance when assembled manually from separate components. We selected top performing conditions from Batch#1 (ePURE1 from round 1 and ePURE2 from round 2) and from Batch#2 (ePURE3 from round 5 and ePURE4 from round 8), and a low-performing condition from Batch#1 (badPURE from round 2) as a control (**Figure 4A**). The maximum mEYFP yield and apparent translation rate were extracted from the fluorescence kinetics. All ePURE compositions outperformed the corresponding REF for both batches of reagents and at both 0.1 nM and 2 nM DNA (**Figure 4**), which is consistent with the performances found with Echo assembly of PURE reactions. While the commercial PURE*frex* 2.0 kit assembled from Solutions I, II and III directly provided by the supplier performs better than ePURE at 2 nM *meyfp* (the recommended concentration of DNA), ePURE1 and ePURE2 are better compositions at 0.1 nM DNA.

**Figure 4:**
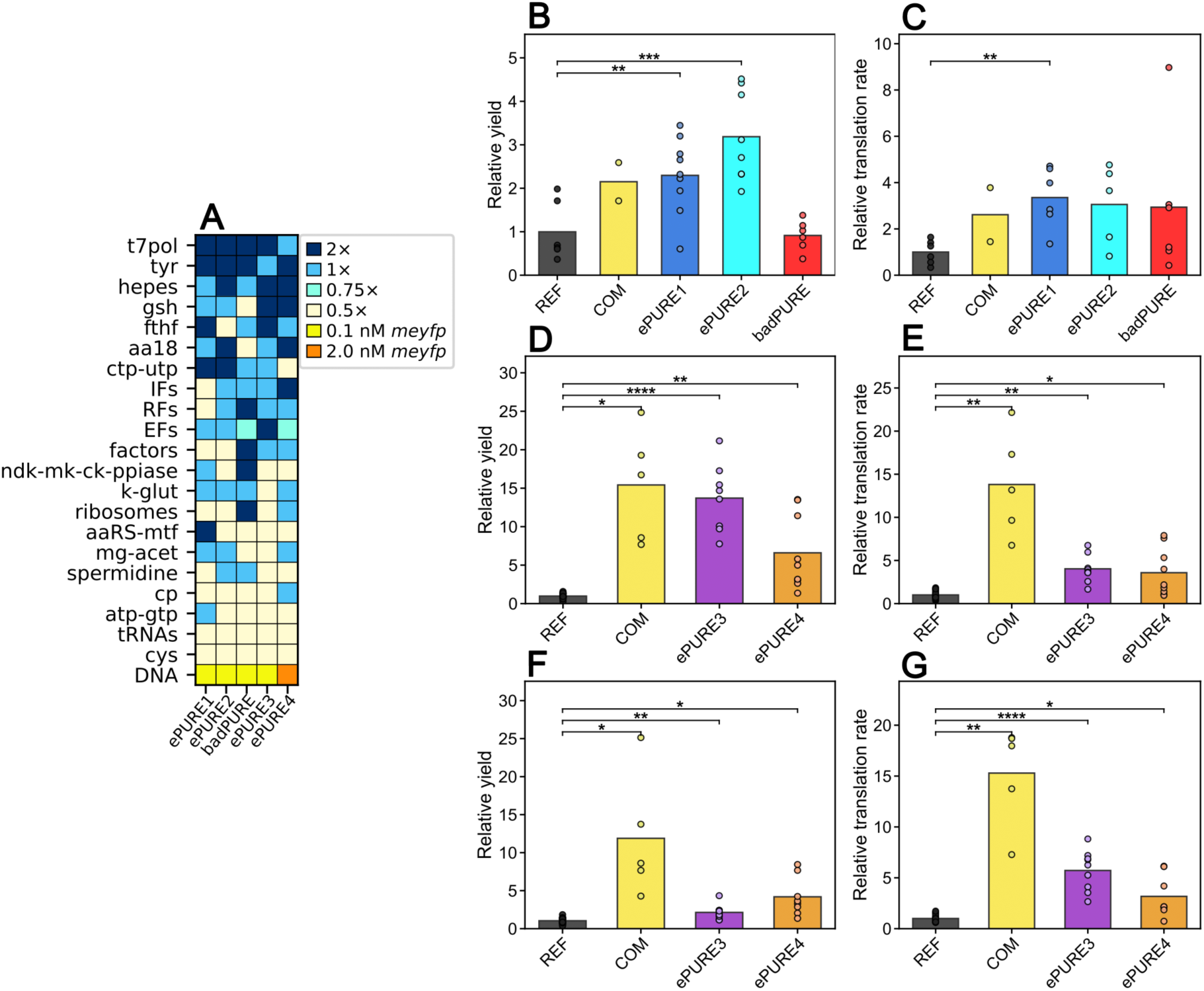
Manual assembly validates enhanced PURE (ePURE) compositions. **(A)** Heatmap showing the relative concentrations of PURE components in different formulations. ePURE1 and ePURE2 were top-performing conditions from rounds 1 and 2 (Batch#1), respectively, badPURE was a low-performing condition from round 2, and ePURE3 and ePURE4 were optimized compositions from rounds 5 and 8 (Batch#2), respectively. Color coding indicates concentration relative to the REF condition. **(B-G)** Bars represent mean values with individual replicate data points appended as circles (n = 2 to 18). Welch’s unpaired (independent) two-sample *t*-test was performed. Statistical significance: \**p* < 0.05, \*\**p* < 0.01, \*\*\**p* < 0.001, \*\*\*\**p* < 0.0001. Relative yield (B) and relative translation rate (C) of mEYFP synthesis at 0.1 nM DNA for Batch#1 formulations. Relative yield (D) and relative translation rate (E) of mEYFP synthesis at 0.1 nM DNA for Batch#2 formulations. Relative yield (F) and relative translation rate (G) of mEYFP synthesis at 2 nM DNA for Batch#2 formulations.

### Active learning applied to the expression of a multicistronic plasmid reveals gene-specific improvement

Having established that active learning can generate new PURE compositions that enhance expression of single-gene reporters, we next sought to maximize proteome-scale expression of a multicistronic DNA containing genes under control of identical regulatory sequences (T7 promoter, ribosome binding site, leader sequence, and transcription terminator) (**Figure 5A**). Specifically, we asked whether such an optimization approach relying on a fluorescent reporter gene would lead to uniform enhancement of all encoded proteins, or instead results in non-uniform effects across genes, leading to PURE composition-dependent protein expression patterns (**Figure 5B**).

**Figure 5:**
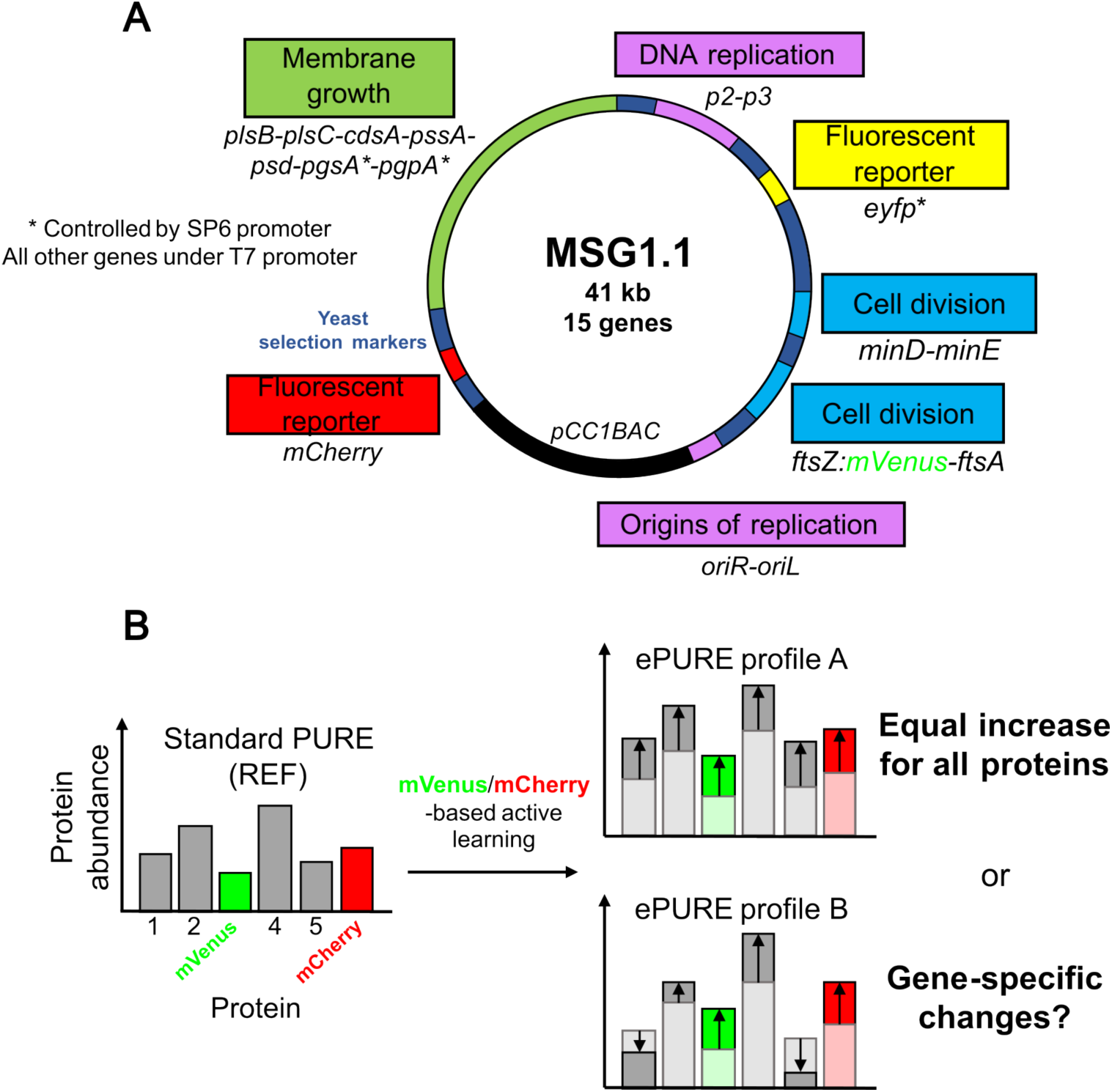
A 15-gene synthetic chromosome for expression in PURE system. (**A**) Schematic representation of the multicistronic plasmid MSG1.1^30^ (41 kb) encoding 15 genes involved in membrane synthesis, DNA replication, and cell division, together with three fluorescent reporter genes. Genes under control of the T7 promoter (pT7) are expressed in PURE system, including two fluorescent reporters (*mVenus* fused to *ftsZ* and *mCherry*) and eleven functional genes: *ftsZ*, *ftsA*, *minD*, *minE*, *plsB*, *plsC*, *cdsA*, *pssA*, *psd, p2, p3*. Additional transcriptional units under SP6 (*eYFP*, *pgsA* and *pgpA*) or yeast promoters are present on the plasmid but are not expressed in PURE. Identical regulatory architectures (T7 promoter, ribosome binding site, and T7 terminator) are used for PURE-expressed genes, except for *p2* that has a double VSV terminator. (**B**) Conceptual representation of possible scenarios for protein abundance profiles following multi-gene expression optimization guided by mVenus and mCherry fluorescence. The left panel depicts the baseline proteomic output after expression of MSG1.1 with the standard PURE system (REF). Active-learning optimization targeting increased mVenus and mCherry yields could lead to two distinct protein profiles. In profile A, increased reporter expression is accompanied by a uniform upward shift in protein yields, reflecting a global enhancement of gene expression capacity that similarly benefits each synthesized protein. In profile B, reporter expression increases while other proteins exhibit heterogeneous responses, indicating gene-specific optimization.

We used MSG1.1, a 41-kb synthetic chromosome encoding 15 genes involved in membrane growth, DNA replication, and cell division, together with three fluorescent reporters^30^ (**Figure 5A**). A subset of genes, including two reporter genes (*mVenus* and *mCherry*), are transcribed by T7 RNA polymerase, whereas other genes, including yeast selection markers, rely on distinct promoters (SP6 and yeast promoters) and are therefore not expressed in PURE system. We applied the active learning workflow to MSG1.1 expression at two DNA concentrations: 0.1 nM and 1 nM (**Figure 6**).

**Figure 6:**
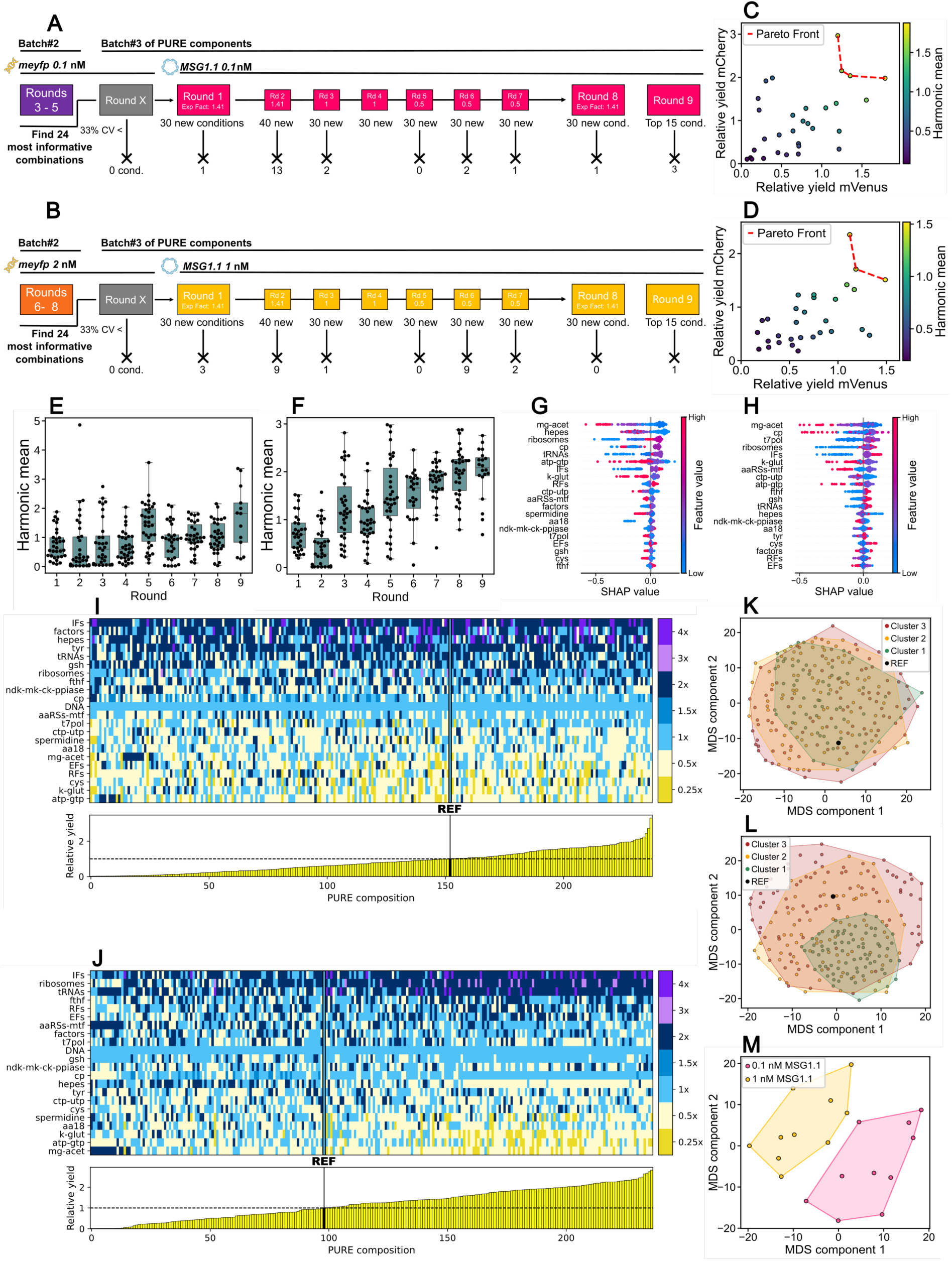
Active learning-based optimization of MSG1.1 expression at 0.1 nM and 1 nM DNA. (A-B) Schematic overview of the optimization strategy. Informative conditions from rounds 3–5 (0.1 nM *meyfp*) and 6–8 (2 nM *meyfp*) seeded MSG1.1 optimization campaigns at 0.1 nM (A) and 1 nM DNA (B), respectively. Each campaign proceeded through eight rounds (∼30 conditions per round). **(C-D)** Scatter plot of relative mVenus versus mCherry yields for individual MSG1.1 optimization conditions at 0.1 nM (C) and at 1 nM DNA (D). Colors indicate the harmonic mean of both reporter yields, and the dashed line marks the Pareto front for multi-objective optimization. **(E-F)** Yield evolution across rounds, quantified as the harmonic mean of mVenus and mCherry fluorescence, at 0.1 nM (E) and 1 nM MSG1.1 (F). Box plots show median (center line), interquartile range (IQR, box), and 1.5× IQR (whiskers), with individual data points overlaid. **(G-H)** Shapley values (SHAP) beeswarm plot showing components most influencing yield prediction, at 0.1 nM (G) and 1 nM MSG1.1 (H). **(I-J)** Kymographs showing relative component concentrations across all rounds, ordered by performance at 0.1 nM (I) and 1 nM DNA (J). Colors represent different component concentrations relative to the REF composition. Subplots display the relative yield for all compositions tested. **(K-L)** MDS representation (Manhattan distances) of all conditions with k-means clustering (k = 3) for 0.1 nM (K) and 1 nM DNA (L). Colors represent clusters ranked by harmonic mean, with cluster 1 in green containing top performing conditions. **(M)** MDS comparing the top 10 conditions from each MSG1.1 concentration, highlighting distinct parameter space regions corresponding to their respective optimal regimes.

To leverage prior knowledge, we transferred 24 most informative conditions from earlier mEYFP campaigns: rounds 3–5 (0.1 nM DNA) and rounds 6–8 (2 nM DNA) informed the initial rounds of MSG1.1 at 0.1 nM and 1 nM, respectively (**Figure 6A and B**). Technical constraints required manual distribution of MSG1.1 as the DNA stock solution exceeded Echo’s handling capacity of 30 ng/µL, while all the other components were dispensed with the robot. In total, eight rounds of optimization were performed, testing ∼30 conditions per round in duplicates (45 in round 2; **Figure 6A,B**). In round 4, one technical replicate for each condition exhibited poor or no expression at all, so only the functioning reactions were kept, and no conditions were discarded based on CV% (**Figure 6A,B**). To optimize MSG1.1 expression, we treated the simultaneous yields of mVenus and mCherry as a multi-objective problem. We defined the objective function for active learning as the harmonic mean of both reporters. This metric favors balanced enhancement and penalizes conditions that improve only one. Pareto analysis in round 1 allowed us to visually identify top performing compositions where neither reporter’s yield is severely compromised (**Figure 6C,D**). This approach ensured harmonic was a valid metric for selection of PURE compositions for MSG1.1 expression (**Figure S7**).

At 0.1 nM DNA, the yields of mVenus and mCherry increased to reach relative harmonic mean values around ∼2 (**Figures 6E and S8**). At 1 nM DNA, gradual improvement was more obvious with harmonic mean values up to ∼3 (**Figures 6F and S8**). For rounds 6–8, concentration ranges of several components were expanded based on their distributions in earlier high-performing conditions to unlock new improved conditions outside the initial parameter space (**Table S1**). Hepes, Ribosomes, IFs, factors and tRNAs were supplied at higher levels (up to 3–4×), while several substrates and cofactors (k-glut, mg-acet, spermidine, atp-gtp, aa18, cys, EFs, RFs and t7pol) were added at lower concentrations (0.25×). Some components (e.g., gsh, cp) were fixed at 1× given that their concentrations were mostly fixed by the model as well as to reduce the parameter space.

To evaluate model predictability for MSG1.1 expression, we applied the nested 5-fold cross-validation framework to datasets at 0.1 nM and 1 nM DNA (**Figure S9**). The number of experimental data points was substantially larger than in previous campaigns (238 and 237 data points), improving statistical power and reducing variance across outer folds. At 0.1 nM DNA, predictive performance of the model was modest with R² = 0.22 ± 0.06 (**Figure S9A**). The persistence of low predictability despite the expanded dataset indicates that harmonic mean variability is difficult to infer from composition alone in these conditions. Nevertheless, feature importance analyses were internally consistent and identified mg-acet as the dominant contributor, followed by cp, t7pol and ribosomes (**Figures 6G and S9B**). In contrast, at 1 nM DNA, model performance improved drastically, with R² = 0.62 ± 0.06 (**Figure S9C**). Feature importance rankings became more stable and structured across gain-, permutation-, and SHAP-based analyses, with hepes, mg-acet, ribosomes, and cp emerging as the principal determinants of yield (**Figures 6H and S9D**). Together, these results indicate that the improved predictability observed for MSG1.1 at 1 nM reflects both a transition to a regime where differences between compositions are efficiently captured and an increased robustness afforded by the larger dataset, whereas the low-DNA condition remains dominated by noisy, weakly predictable effects.

Kymograph analysis of all MSG1.1 expression conditions reveals distinct compositional organization at 0.1 nM and 1 nM DNA (**Figure 6I,J**). At 0.1 nM DNA, high-yielding conditions show no clear compositional signature, with component concentrations varying substantially across conditions (**Figure 6I**). In contrast, at 1 nM DNA, the kymograph shows a clear compositional pattern associated with improved performance (**Figure 6J**). High-performing compositions consistently converge toward increased IFs, ribosomes and tRNA concentrations, while mg-acet, atp-gtp, k-glut and hepes are reduced. Component distributions are markedly narrower in this regime, indicating stronger compositional constraints associated with higher yields.

These trends are also reflected in the corresponding MDS representations (**Figure 6K,L**). At 0.1 nM DNA, all three clusters of conditions with different performance levels occupy highly overlapping regions of the MDS space (**Figure 6K**). In contrast, at 1 nM DNA, high-performing conditions collapse into a compact cluster (cluster 1; 1.688 ≤ relative yield ≤ 2.817) that is clearly distinct from lower-performing compositions (**Figure 6L**). Direct comparison of the ten highest-performing conditions from the 0.1 nM and 1 nM MSG1.1 campaigns shows that they occupy largely non-overlapping regions of the MDS space (**Figure 6M**). This separation indicates that optimal compositional regimes differ between low and high concentrations of MSG1.1.

To quantify relationships among PURE components and their importance to protein expression output, we computed pairwise Spearman correlation matrices with relative yield, at low (0.1 nM) and high (1 nM) MSG1.1 concentrations (**Figure 7**). This analysis captures both component–component covariation across the tested compositions and monotonic relationships between individual components and yield. At 0.1 nM DNA, the correlation matrix revealed generally weak pairwise correlations among components (**Figure 7A**), as well as moderate correlation between the individual components and yield (**Figure 7B**). At 1 nM DNA, the correlation structure was more noticeable with higher values of absolute Spearman correlation coefficients compared to the low-DNA condition (**Figure 7C**). Strong positive correlations appeared between k-glut and both hepes and mg-acet, while mg-acet also showed a strong positive correlation with hepes. Spermidine and k-glut exhibited a moderate positive correlation. In contrast, strong negative correlations were observed between ribosomes and hepes, k-glut, and mg-acet. Additional moderate negative correlations were detected between IFs and hepes, as well as between tRNAs and K-glut (**Figure 7D**), where relative yield increased more systematically with higher ribosome, IF, and tRNA concentrations, and decreased with increasing hepes, mg-acet, and k-glut.

**Figure 7.**
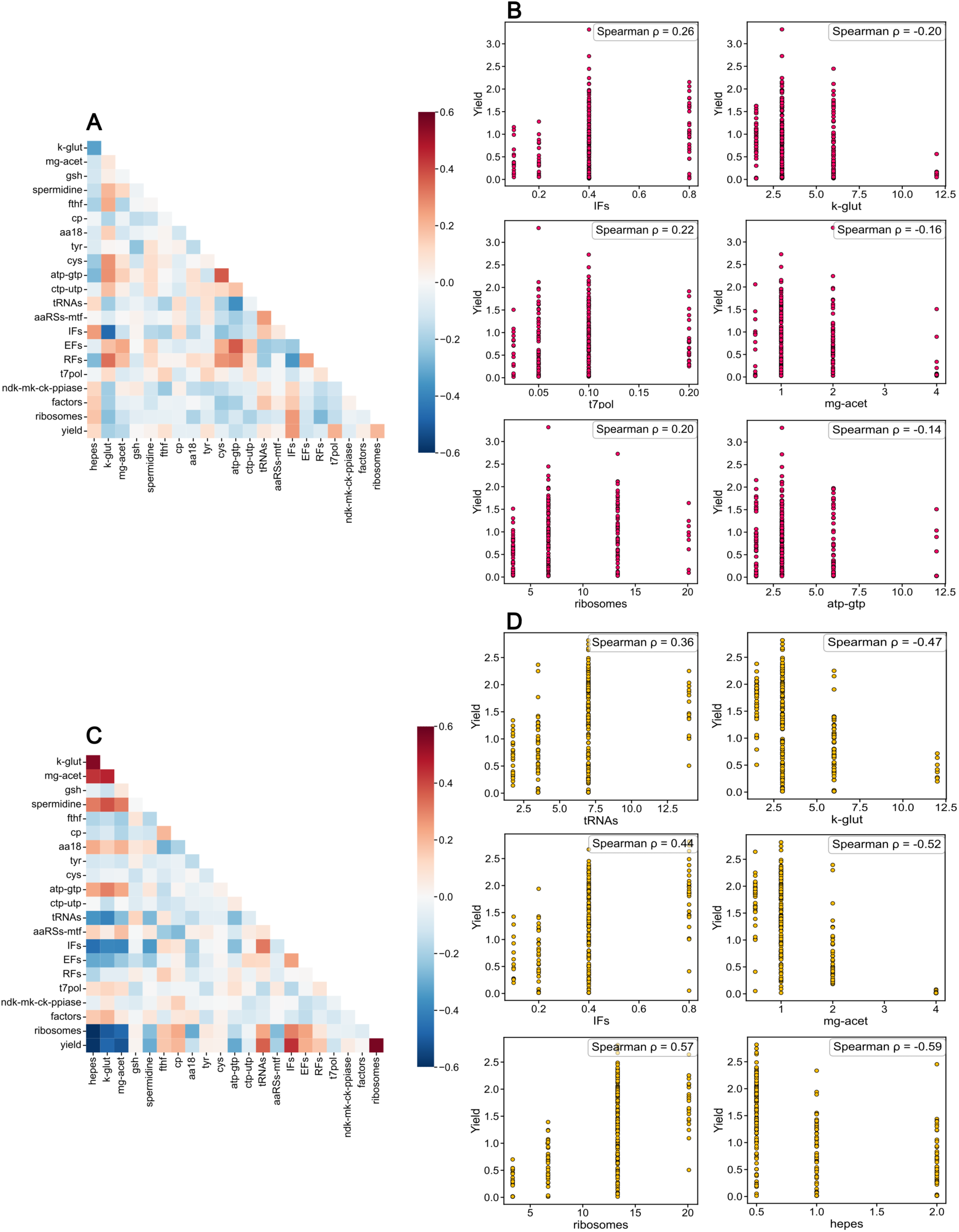
Correlation analysis of PURE component concentrations and expression yield at low and high MSG1.1 concentrations. Spearman correlation matrix between normalized PURE component concentrations and relative yield for experiments performed at 0.1 nM **(A)** and 1 nM **(C)** MSG1.1. Colors indicate the direction and magnitude of correlations (red, positive; blue, negative). Scatter plots showing the relationship between relative yield and selected PURE components: IFs, t7pol, ribosomes, k-glut, mg-acet, and atp-gtp at 0.1 nM **(B)**, and ribosomes, IFs, tRNAs, hepes, mg-acet, and k-glut at 1 nM **(D)** MSG1.1. Each point represents one PURE composition. Spearman correlation coefficients (ρ) are reported in each panel.

Finally, to test whether optimization that was guided by the fluorescent reporters also improved expression of the other genes of MSG1.1, we quantified individual protein abundances in the best-performing PURE compositions by mass spectrometry (**Figure 8A,B**). We selected ePURE5 (0.1 nM DNA) and ePURE6 (1 nM DNA) from round 3, ePURE7 (0.1 nM DNA) and ePURE8 (1 nM DNA) from round 7, and ePURE9–12 (0.1 nM DNA) and ePURE13–16 (1 nM DNA) from round 9. Although optimization effects were generally weaker at 1 nM DNA than 0.1 nM, the rough protein expression profiles indicate high variability in the relative abundances, with some proteins being overexpressed (e.g., mCherry and to a less extent mVenus and CdsA) and some underexpressed (MinD, PlsB, DNA pol.) compared to the REF composition. This is remarkable, given that all the genes have similar regulatory sequences. Regarding the two proposed optimization outcomes illustrated in **Figure 5B**, mass spectrometry data reveal that the proteomic profiles with ePURE are clearly more in line with a gene-specific optimization scenario. These results suggest that improving protein synthesis yield from individual genes or a subset of genes within a multicistronic DNA is feasible but would require a specific readout of the target protein(s) for steering the active learning.

**Figure 8:**
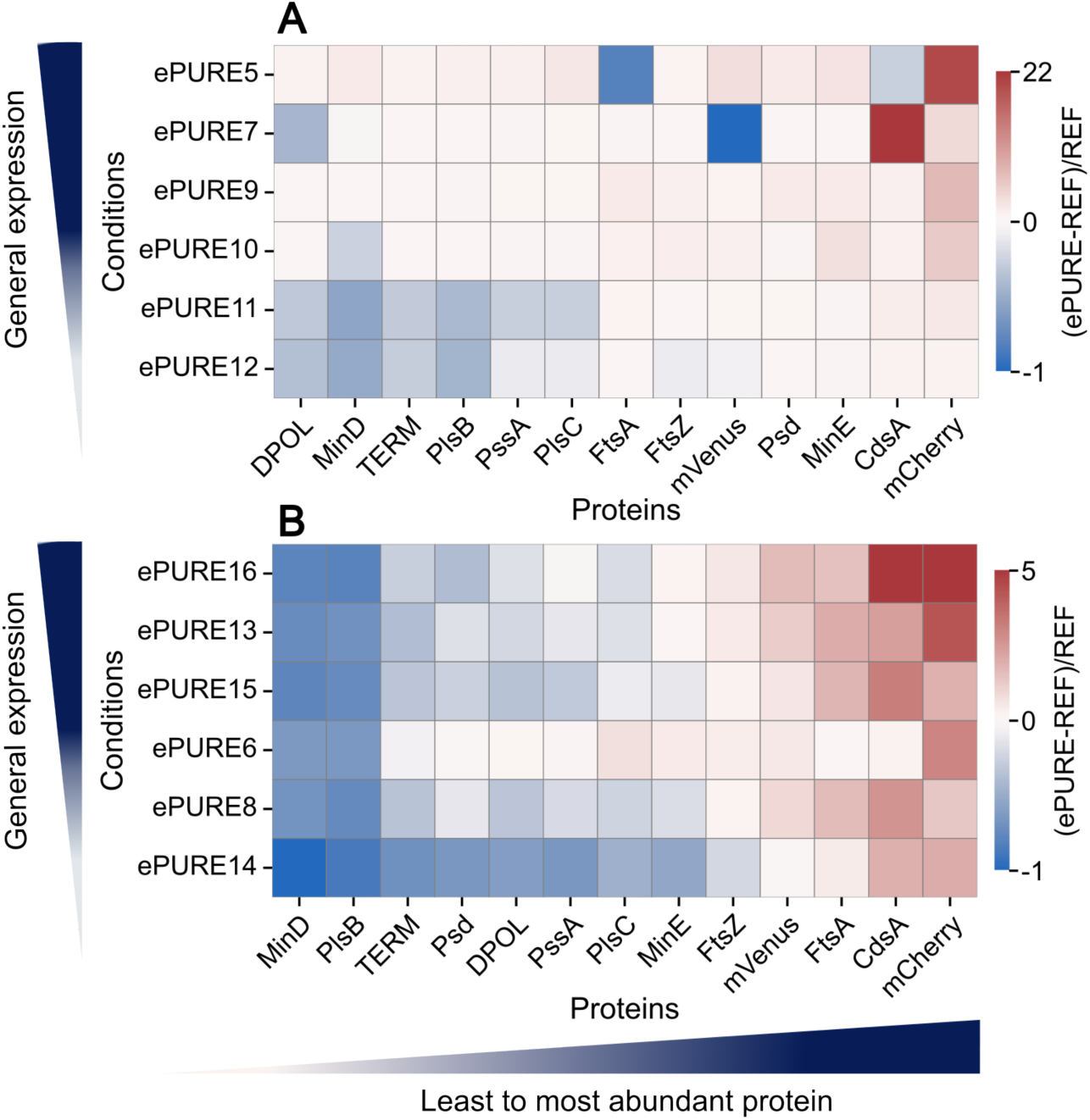
Proteomic analysis reveals no general optimization of MSG1.1 expression across the encoded genes. Heatmaps of relative protein abundances resulting from the expression of MSG1.1 at 0.1 nM **(A)** and 1 nM **(B)** in different ePURE compositions. Color coding is the normalized raw protein abundance after expression in ePURE conditions ranked from top to bottom by summing the individual protein abundances (global expression capacity).

## DISCUSSION

In this study, we developed an active-learning-guided approach for optimizing the composition of PURE system^4,5^, demonstrating that data-driven exploration can efficiently navigate the large and highly multidimensional parameter space of a CFE system^10,21,23^. Despite the combinatorial scale of the problem, higher-performing compositions were identified within only a few iterative rounds, highlighting the structured composition-efficiency landscape of PURE system. The emergence of narrow regions of high-performing conditions in MDS space further indicates that the algorithm rapidly converges toward relevant compositional configurations, in contrast to what would be expected from random or unguided exploration. Notably, the degree of yield improvement and relationship between the molecular composition and CFE efficiency depends on the concentration of DNA and on the batch of purified components.

This study explored a rather small range of concentrations (initially 0.5× to 2×), which may have constrained the discovery of more extreme, yet potentially beneficial compositions. Practical limitations also restricted throughput for large plasmids like MSG1.1, as DNA supply required manual pipetting. Moreover, some degree of batch-to-batch variability was observed, which could pose challenges for reproducibility^31^. Despite these variations, the active-learning workflow consistently identified higher-yielding compositions within each batch. Echo- and manually assembled reactions performed comparably, indicating that the optimized conditions are independent of the assembly method. However, custom ePURE mixtures achieved lower absolute yield compared to the commercial PURE*frex* 2.0 kit in most conditions tested (**Figure 4**). This finding underlines adaptive optimization not only as a path to surpassing commercial formulations, but as a means for laboratories to dynamically recalibrate tailored PURE preparations to accommodate batch specificities.

Across rounds and experimental conditions, a strong positive correlation was found between translation rate and final protein yield, underscoring the central role of translational efficiency in determining PURE productivity^19,32^. This relationship suggests that improving translational efficiency, for instance by enhancing the fraction of active ribosomes^33,34^, the efficacy of translation initiation and ribosome processivity^35^, or ATP regeneration^18^, may represent a primary route to enhancing overall protein output. The systematic enrichment or depletion of specific substrates and cofactors in high-yielding compositions further indicates that the REF formulation contains latent stoichiometric imbalances. In particular, the depletion of cp, atp-gtp, mg-acet, and tRNAs in many top-performing conditions suggests that their standard concentrations may exceed the optimal balance required for efficient energy turnover or translation. This observation is especially encouraging for efforts toward self-regenerating PURE systems^33,36–38^, as it implies that reducing the concentrations of certain components may be feasible without compromising performance. Conversely, the repeated enrichment of tyr, t7pol, IFs, tRNAs or ribosomes suggests that the amount of these components could be increased for a wider array of expressed DNA. In this way, active learning works not only as an optimization strategy but also as a diagnostic tool, capable of revealing which components require tweaking to reach a target function.

It is important to note, however, that these observations remain context-specific, as not all trends are observed across every optimization campaign. For instance, a major outcome of this work is the identification of distinct limiting regimes governing PURE performance as a function of DNA template concentration. While prior studies suggested that T7 RNA polymerase activity is not a limiting factor in PURE system^17^, we found that it becomes a bottleneck at low DNA loads (0.1 nM *meyfp*), where the concentration of t7pol strongly influences protein yield (**Figure 2H**). At higher DNA concentrations, the system shifted to a qualitatively different regime most sensitive to translation initiation, where high concentrations of IFs favor high yield. This context-dependency is particularly relevant given the molecular diversity among existing PURE formulations, which range from commercial kits with undisclosed compositions like PURExpress and PURE*frex* to laboratory-prepared concoctions such as PURE3^39^ and One-Pot PURE^40^. Because different PURE system variants possess different component concentrations and levels of purity, our results may not readily apply to other CFE systems. However, the presented workflow can be seamlessly adapted to optimize these alternate PURE variants by simply adjusting the components and parameter space accordingly. Together, these results indicate that PURE system does not operate under a single universal limiting factor; rather, its constraints depend strongly on template load and biochemical context.

Comparisons across DNA concentrations and component batches revealed partial convergence of high-performing compositions into a shared, compact region of the MDS landscape, representing a potential ePURE subspace associated with generally favorable biochemical conditions. However, the limited overlap between optimal compositions across DNA loads argues against the existence of a ‘universal’ PURE recipe. Instead, the data support a context-dependent landscape in which optimality depends on template concentration, template architecture (and possibly genetic organization), and component batch characteristics, reinforcing the idea of adaptable PURE formulations tailored to specific applications rather than a single fixed composition^10^.

In parallel with these biochemical insights, the behavior of the active learning models themselves provides additional information about the structure of the PURE optimization landscape^21–23^. Across campaigns, model performance varied strongly with DNA concentration, template complexity, and dataset size, reflecting intrinsic differences in how tightly molecular composition constrains functional outcome. At low DNA concentrations and in early single-gene campaigns, predictive power remained modest, with models capturing coarse trends and relative ranking rather than absolute yields. In these regimes, predicted values were systematically compressed toward intermediate outputs, suggesting that yield variability is influenced by stochastic or weakly structured effects not fully encoded by component concentrations alone^41^. Nevertheless, feature-importance analyses remained internally consistent and reproducible, indicating that even when predictive accuracy is limited, the models extract biologically meaningful patterns^42–45^.

By contrast, at higher DNA concentrations and in later campaigns, most notably at 2 nM *meyfp* and 1 nM of MSG1.1, model performance improved substantially, with a larger fraction of variance explained and clearer separation between low- and high-performing conditions. This improvement coincided with tighter clustering in MDS space and more pronounced kymograph patterns, indicating that PURE behavior becomes more deterministic and composition-driven in these regimes. Importantly, the increase in predictive power was not solely a consequence of larger datasets but rather reflected a transition toward a functional state governed by fewer, more dominant constraints, primarily linked to translation initiation, as well as ribosome and tRNA availability. Model performance therefore serves as a quantitative readout of the structure of the molecular landscape (composition space): high predictability corresponds to well-defined processes limiting yield, whereas low predictability reflects diffuse or shifting constraints in the PURE molecular composition. The increased predictive power of the model further highlights the advantage of active learning over random or intuition-driven exploration. While random sampling was sufficient to identify occasional high-performing outliers in early rounds, these conditions were sparse and suboptimal. In contrast, the active-learning strategy rapidly enriched for high-yielding compositions, generating compact clusters of related formulations in MDS space. This behavior indicates that the algorithm does not merely identify isolated optima, but learns the local structure of favorable regions, enabling systematic refinement across rounds^46^.

Extending the workflow to the expression of the 41-kb synthetic chromosome MSG1.1^30^ further demonstrated the limits of composition-only optimization. While the production of mCherry and mVenus increased under optimized conditions, several genes exhibited weaker or unpredictable responses despite sharing identical regulatory sequences. Mass spectrometry revealed that improvements based on a target protein were not uniformly transmitted across the proteome. This observation suggests that, for large multicistronic templates, limiting factors are not bounded to global resource availability but encompass sequence-encoded features such as transcriptional context, mRNA structure, codon usage, as well as complete protein translation paired with folding efficiency^47^. Thus, although system-level optimization can enhance expression of specific genes, it does not uniformly increase protein output across a large gene set and cannot compensate for sequence-encoded constraints. Future optimization strategies will therefore need to integrate compositional tuning with template-level design principles.

The space of possible PURE reaction compositions is still large. Increasing throughput and iterations and expanding concentration ranges of the components will accelerate the discovery of PURE formulations with higher robustness and protein synthesis capacity. Moreover, incorporating multi-objective optimization criteria such as reagents cost^48^, translation rate, production lifetime, genome-wide protein abundance profile^49^, and integrating sequence-level predictors^50^ represent even more powerful optimization strategies, especially when implemented in a closed-loop autonomous workflow^51^. Finally, generalization of the results to other reaction formats like microcompartments (liposomes) still needs to be shown for the development of PURE-based synthetic cells.

## MATERIALS AND METHODS

### Plasmids and DNA preparation

Linear DNA was used for *in vitro* expression of *meyfp* (monomeric Enhanced Yellow Fluorescent Protein) and *lacZ* (with gene product β-galactosidase). Templates were obtained through polymerase chain reaction (PCR) using plasmid DNA G76 or pT003. G76 is a pUC57-derived plasmid containing *meyfp* expression cassette under the control of a T7 promoter. It was synthesized and codon-optimized for *E. coli* by Eurogentech^52^. pT003 is also derived from pUC57 and contains a *lacZ* expression cassette under a T7 promoter^53^. PCR reactions were performed using either Phusion or Q5 High-Fidelity DNA polymerases (NEB) and the products were purified using the NucleoSpin PCR clean-up kit (Macherey-Nagel) or the QIAquick PCR Purification Kit (Qiagen). Quantification was performed using a Thermo Scientific NanoDrop ND 1000 Spectrophotometer.

Plasmid pUDF006-MSG1.1 (referred to as MSG1.1) was assembled as described in ref. ^30^ and was used here in its circular form for *in vitro* expression. MSG1.1 was isolated from *E. coli* EPI300 cells using the NucleoBond Xtra Midi kit (Macherey-Nagel). To avoid mixed or recombined plasmid populations, colonies were first isolated on LB agar supplemented with chloramphenicol (12.5 µg/mL). A glycerol stock of MSG1.1 was streaked to obtain single colonies, which were screened by colony PCR using DreamTaq DNA polymerase (Thermo Fisher Scientific) to verify the presence of the mRuby2, CEN6/ARS4, and pCC1BAC backbone sequences. Verified colonies were used to inoculate 10 mL overnight cultures, which were subsequently scaled up to 300 mL LB + chloramphenicol (12.5 µg/mL). High-copy-number induction of the pCC1BAC origin was performed prior to plasmid isolation to maximize DNA yield. Plasmid preparations were checked by agarose gel electrophoresis (0.6%) after linearization with PmeI to confirm plasmid size and homogeneity. DNA concentration was measured using both NanoDrop and Qubit fluorometric quantification (Thermo Fisher Scientific). Correct midipreps were pooled, aliquoted, and stored at 4 °C to prevent mechanical shearing. Plasmid identity and integrity were further confirmed by Plasmidsaurus Inc. whole-plasmid sequencing. All plasmids and primers are described in **Tables S2 and S3.**

### PURE components handling

PURE components were prepared by GeneFrontier Corporation (Japan) and stored at −80°C. Thawing on ice was performed before use and the remaining components were flash-frozen using liquid nitrogen. For each active learning round, fresh components were used that were never thawed before. Stock solutions of components in Solution I of PURE*frex* 2.0 (hepes, k-glut, mg-acet, gsh, spermidine, fthf, cp, cys, ctp-utp) were diluted with RNase-free water and handled at room temperature to avoid precipitation. Other Solution I components (aa18, tyr, atp-gtp, tRNAs) were used directly from the supplied tubes. Components in Solution II (aaRS-mtf, Ifs, EFs, RFs, t7pol, ndk-mk-ck-ppiase, factors) were diluted on ice using a dilution buffer (HEPES-KOH (pH 7.6) 20 mM, Potassium Acetate 100 mM, 3-Mercapto-1,2-propanediol 7 mM, Glycerol 30%). Ribosomes (Solution III) were used without further dilution.

### Echo dispensing and PURE reaction assembly

Reactions were assembled using an Echo 525 acoustic liquid handler (Beckman Coulter Life Sciences). Ready-to-use components (individual or pre-mixed PURE components) were loaded onto a 384PP+ source plate (Beckman Coulter). Calibration tests were performed for each component to identify the optimal liquid class among the five calibration options available for the 384PP+ plate. Eight 200-nL droplets were dispensed on a filmed 384-well black, clear-bottom Greiner plate (Ref. 781906). Assessment was based on visual inspection of the droplet size and quality (shape/satellite droplets) and on the volumes remaining in the source well and dispensed with Echo as measured with surveys performed before each dispensing command. Before each Echo experiment, the source plate was loaded at room temperature, centrifuged briefly and gently mixed to remove air bubbles before dispensing.

Echo transfer scripts in the form of .csv files were generated using a customized METIS optimization workflow^22^. The “Transform Volumes.csv to Echo Liquid Handler Compatible Input” module was modified to automate experimental setup, destination plate layout generation, including calculation of required total source volumes and source plate layout generation. Dispensing order was selected to fit GeneFrontier’s recommendations. Each 11 µL reaction was assembled in 384-well black Greiner plates using acoustic liquid handling, with all component volumes dispensed in 25 nL increments. Each reaction included either 100 pM *meyfp* + 10 pM *lacZ* templates or 2 nM *meyfp* + 0.1 nM *lacZ* by dispensing a fixed 0.4 µL DNA solution. MSG1.1 plasmid was added manually by pipetting 0.4 µL to reach final concentrations of 0.1 nM or 1 nM.

All conditions were assembled in technical duplicates or triplicates and the measurement plate was sealed with an optically clear adhesive film prior to incubation at 37 °C.

### Manual assembly of PURE reactions

Selected reactions, including conditions optimized in early rounds, were assembled manually for validation of the protein synthesis yields and rates found with Echo-dispensed components. Solution I components were mixed at room temperature to avoid precipitation, whereas Solution II components were combined on ice. Master mixes were assembled and 11 µL per reaction (3 technical replicates) was transferred into a 384-well Greiner microplate (Ref. 781906) prior to fluorescence kinetic measurement. PURE*frex* 2.0 reactions with the commercial kit were prepared following the manufacturer’s instructions.

### Fluorescence and absorbance kinetics

The microplate was sealed with an optical adhesive cover from Applied Biosystems (Ref. 4360954) and fluorescence kinetics of mEYFP expression were monitored using a CLARIOstar microplate reader (BMG Labtech) at 37 °C. Fluorescence was measured every 10 min for 16 h using 497 ± 15 nm excitation and 540 ± 20 nm emission filters (bottom read, 20 flashes per well). At the end of the reaction, chlorophenol red-β-D-galactopyranoside (CPRG, 0.5 mg mL⁻¹ final) was added manually or via Echo to measure β-galactosidase activity. Enzymatic conversion of CPRG to CPR was monitored at 575 nm every 3 min for up to 4 h at 37 °C. For MSG1.1 expression, fluorescence kinetics of mVenus and mCherry were monitored simultaneously. Fluorescence was measured every 5 min for 16 h using 497 ± 15 nm excitation and 540 ± 20 nm emission filters for mVenus, and 570 ± 15 nm excitation and 620 ± 20 nm emission filters for mCherry (bottom read, 20 flashes per well). Raw data were processed using CLARIOstar MARS software and exported to Excel format.

### Fluorescence and absorbance data analysis

Raw fluorescence data were processed using a custom Python script. End-point mean fluorescence values, standard deviations and coefficients of variation (CV %) were computed across technical replicates. Conditions with CV > 33% were excluded from downstream analysis. Relative yields were obtained for each condition as the maximum value of mean fluorescence after 16 h divided by the maximum value from the mean of REF. To reduce the impact of noise in fluorescence kinetics in MSG1.1 experiments, maximum fluorescence values for mVenus and mCherry were estimated using a sliding-window (window size = 5 consecutive data points). The mean fluorescence was calculated for all consecutive five-point windows across the time course, and the highest mean value was retained.

Relative yields were then calculated as the ratio between the maximum window-averaged fluorescence of each condition and that of the REF condition. A phenomenological model^54^ was used to extract translation rate and plateau time, based on the following sigmoid equation:

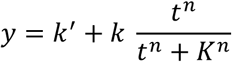

where *t* is the time in hours, *y* is the fluorescence (RFU value) at a given time point, and *k’*, *k*, *K* and *n* are fit parameters. The apparent translation rate is defined as the steepness at time *t* = *K* and is expressed as:

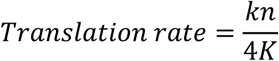

Absorbance data were analyzed using a Python script. Mean absorbance values across technical replicates were calculated and the steepest slope of each curve was determined using a sliding-window as described above, and relative slopes were obtained by normalizing each value to the steepest slope of the reference (REF) condition.

### METIS workflow for active learning

METIS Optimization Notebook^22^ was used and adapted through Google Collab (Python 3) and executed with the following experimental parameters: each round included between 25 to 55 new combinations (*m* variable), the minimum droplet volume was set to 25 nL, and the total reaction volume was 11 µL. In the METIS *concentration_limits* dictionary, each PURE component was defined as a feature with three discrete values representing the three possible relative concentrations compared to the REF condition. The *Conc_Stock* values indicated dilution factors (100 = undiluted stock, 33.3 = threefold dilution). The *specials* dictionary included the REF composition and selected high-performing conditions that were re-tested in subsequent rounds. Every round, *Concentrations_X.csv* files were generated containing all the conditions to be tested with the concentration of each component. This file was then transformed into a *Volumes_X.csv* file based on experimental parameters. For the first round (Round 0), “Day 1” module was used leading to the generation of random combinations which served as the starting point for the active learning campaigns. During active learning, *Results_X.csv* (with “X” corresponding to the round number) were generated by modifying *Concentrations_X.csv* files with an added “yield” feature that corresponded to the relative yield of mEYFP expression or the relative yield (harmonic mean) of mCherry and mVenus for MSG1.1 expression.

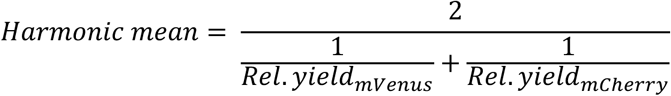

Standard deviation and coefficient of variation (CV) were calculated for each condition, and only those with CV < 33% were selected for learning. To accommodate input files containing a variable number of conditions, the data-formatting cells in METIS were modified, enabling the appending of condition counts exceeding *m*. Active learning and Bayesian optimization were performed as described in ref. ^22^ using XGBoost regressor with RandomCV for learning (see hyperparameters in **Table S4**), 200 iterations with 5 cross validations to reduce overfitting, then the 20 best models were selected based on their mean squared error (MSE). For Bayesian optimization, 10,000 random conditions were generated, and their yield was predicted with 20 XGBoost regressors. These conditions were then ranked based on their Upper Confidence Bound (UCB) expressed as:

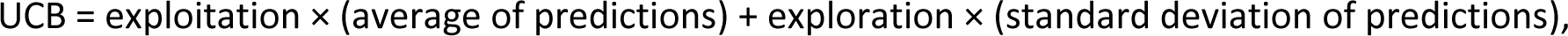

with the term ‘exploitation’ fixed at 1 and ‘exploration’ varying between [1.41; 1; 0.5] depending on rounds. Out of the best ranked conditions, *m* conditions are selected to be tested in the following round. The module *Results as Box Plot* was used for graph generation. The module *Find K Most Informative Combinations* was used to suggest key conditions to be tested in next experiments.

### Computational analysis and visualization

Data processing and visualization were performed using Python 3.11.8 with pandas (v 2.2.1), numpy (v1.26.4), matplotlib (v 3.10.6), and seaborn (v 0.13.2). Data standardization, multidimensional scaling through pairwise Manhattan distance calculations and k-means clustering were performed using scikit-learn (v 1.4.2). Curve fitting, Gaussian smoothing, linear regression, convex hull computation, Welch’s unpaired *t*-test and Spearman nonlinear correlations were performed using scipy (v 1.16.0). For all stochastic procedures, the random seed was set to 42 to ensure reproducibility.

For each active learning campaign, a dataset was created combining conditions from every round. Repeated conditions were curated by averaging the relative yields (and relative rates if present), giving datasets with a unique pair of kinetic parameters for each condition, in the form of *.csv* files. From this, heatmaps (kymographs) were generated, with PURE compositions sorted by relative yield and components ordered by the mean of their relative concentration values (normalized to REF), providing an overview of compositional trends with patterns possibly emerging across the dataset.

To further explore compositional patterns in these high-dimensional datasets and identify similarities between PURE compositions, multidimensional scaling (MDS) was applied using pairwise Manhattan distances as the dissimilarity metric, generating a continuous two-dimensional representation of condition similarity. K-means clustering was then performed on relative yield or harmonic mean values to detect groups of conditions with similar profiles. The optimal number of clusters was determined empirically by minimizing the within-cluster sum of squares and inspecting the silhouette score distribution. Cluster assignments were mapped onto the MDS space to facilitate interpretation of both local and global trends, and convex hulls were computed to visualize the boundaries of different condition groups.

### Machine Learning

To assess how well PURE composition–performance relationships could be captured by machine learning, an independent analysis pipeline was implemented outside the METIS active learning framework using data points obtained across experimental rounds. Gradient-boosted regression models were trained on experimentally measured yields to quantify dataset predictability and evaluate model performance.

All computational analyses were performed using Python 3.11.8 with pandas, numpy, matplotlib, seaborn, scikit-learn, scipy, xgboost (v2.1.2), and shap (v0.46.0). For all stochastic procedures, the random seed was set to 42 to ensure reproducibility. An XGBoost regressor was embedded in a preprocessing pipeline including feature standardization (zero mean, unit variance). Due to a limited dataset and to improve robustness, model performance was evaluated using nested five-fold cross-validation. In the outer loop, data were split into five folds, each held out once for testing. Within each outer training set, an inner five-fold cross-validation was used for hyperparameter optimization using RandomizedSearchCV (200 iterations), minimizing the mean absolute error. The hyperparameter search space and optimization strategy were identical to those used in the METIS active learning workflow (**Table S4**), including learning rate, maximum tree depth, number of estimators, subsampling ratios, regularization strength, minimum child weight, and tree-splitting penalty.

For each outer fold, the best-performing model was evaluated on the held-out test set, and performance was quantified using mean absolute error (MAE), mean squared error (MSE), and coefficient of determination (R²). Final model performance was reported as the average and standard deviation of these metrics across all outer folds. Predictions from all outer folds were aggregated to generate predicted-versus-observed yield plots.

A final XGBoost model was trained on the full dataset, containing all data points previously discarded during cross-validation, using the most frequently selected hyperparameter configuration across outer folds. This model was used for feature importance analysis to identify components most influential on yield predictions by the model. Feature importance was assessed using three complementary methods: XGBoost gain importance, SHAP (SHapley Additive exPlanations) values computed using a randomly sampled background dataset (≤100 samples), and permutation importance based on increases in mean absolute error upon feature shuffling. Each metric was normalized and averaged to obtain a consensus feature importance score. Results were visualized using bar plots and SHAP summary (beeswarm) plots.

### Liquid chromatography mass spectrometry

Echo-assembled PURE reactions containing MSG1.1 were analyzed by liquid chromatography mass spectrometry (LC-MS). Ten microliters of samples were processed for trypsin digestion by addition of ten microliter of trypsin (500 ng) in ammonium bicarbonate (100 mM) to each sample for overnight digestion. The reaction was stopped by adding 1.5 μL 1% trifluoroacetic acid (TFA).

Tryptic peptides were analyzed by Nano-LC coupled to tandem MS using an UltiMate 3000 system (NCS-3500RS Nano/Cap System, Thermo Fisher Scientific) coupled to an Orbitrap Q Exactive Plus mass spectrometer (Thermo Fisher Scientific). Five microliters of sample were injected on a C18 pre-column (300 μm inner diameter × 5 mm, Thermo Fisher Scientific) in a solution consisting of 2% acetonitrile and 0.05% TFA, at a flow rate of 20 μL min^−1^. After 5 min of desalting, the precolumn was switched online with the analytical C18 column (75 μm inner diameter × 50 cm; in-house packed with Reprosil C18) equilibrated in 95% solvent A (5% acetonitrile, 0.2% formic acid) and 5% solvent B (80% acetonitrile, 0.2% formic acid). Peptides were eluted using a 10–50% gradient of solvent B over 105 min at a flow rate of 300 nL min^−1^. The mass spectrometer was operated in data-dependent acquisition mode with the Xcalibur software. MS survey scans were acquired with a resolution of 70,000 and an AGC target of 3×10^6^. The ten most intense ions were selected for fragmentation by high-energy collision-induced dissociation, and the resulting fragments were analyzed at a resolution of 17,500 using an AGC target of 1×10^5^ and a maximum fill time of 50 ms. Dynamic exclusion was used within 30 s to prevent repetitive selection of the same peptide.

Raw MS files were processed with the Mascot software (version 2.7.0) for database search and Proline^55^ for label-free quantitative analysis (version 2.1.2). Data were searched against *E. coli* entries of the UniProtKB protein database release Swiss-Prot 2019_11 (23,135 entries) and a homemade database (built with FASTA sequences of expected proteins mVenus from *Aequorea victoria*, DNAP and TP from *Bacillus* phage φ29 and mCherry from *Anaplasma marginale*). Oxidation of methionine was set as a variable modification. Specificity of trypsin/P digestion was set for cleavage after K or R, and two missed trypsin cleavage sites were allowed. The mass tolerance was set to 10 ppm for the precursor and to 20 mmu in tandem MS mode. Minimum peptide length was set to 7 amino acids and identification results were further validated in Proline by the target decoy approach using a reverse database at both a PSM and protein false-discovery rate of 1%. For relative quantification of proteins across biological replicates and conditions, cross-assignment of peptide ion peaks was enabled within each group with a match time window of 1 min, after alignment of the runs with a tolerance of ± 600 s. Raw abundance values were visualized as bar plots. For relative quantitative comparison between conditions, protein abundances were normalized to the reference formulation (REF). Relative changes were calculated for each protein as (ePURE-REF)/REF, where ePURE denotes the abundance measured in the optimized condition. Data processing and visualization were performed using custom Python scripts. Relative abundance changes were represented as heatmaps generated using the seaborn package, enabling systematic comparison of proteomic profiles between optimized and reference PURE compositions.

## Acknowledgements

We thank Paola Randazzo, Hélène Serrano, and Marion Deville from the Toulouse White Biotechnology platform for assisting with the Echo liquid handler, and Alexandre Stella and Etienne Bancal from the ProteoToul service (IPBS Toulouse) for performing the proteomics analysis. We also thank Alexandre Huyghe for the design of automated scripts and fruitful discussions, Leonardo Morini and Bob van Sluijs from Wilhelm Huck lab for providing us with the initial METIS optimization notebook used in this study, and Pascale Daran Lapujade for her guidance in the preparation of MSG1.1. We are grateful to GeneFrontier for sponsoring our research. Generative AI tools were used for assistance with script and icon generation; all outputs were reviewed and validated by the authors.

## Funding

CC, AS and CD acknowledge financial support from the “BaSyC – Building a Synthetic Cell” Gravitation grant (024.003.019) of the Netherlands Ministry of Education, Culture and Science (OCW) and the Netherlands Organisation for Scientific Research (NWO). CD acknowledges funding from Agence Nationale de la Recherche (ANR-22-CPJ2-0091-01). YBL, MJH and CD also acknowledge funding from the “Bourses de thèse sur enjeux sociétaux” program (EvoCell, 2023) of INSA Toulouse.

## Notes

### Competing Interest Statement

The authors have declared no competing interest.

