## Supplementary Information for "Optimization of PURE system composition using automation and active learning"

**FOR**

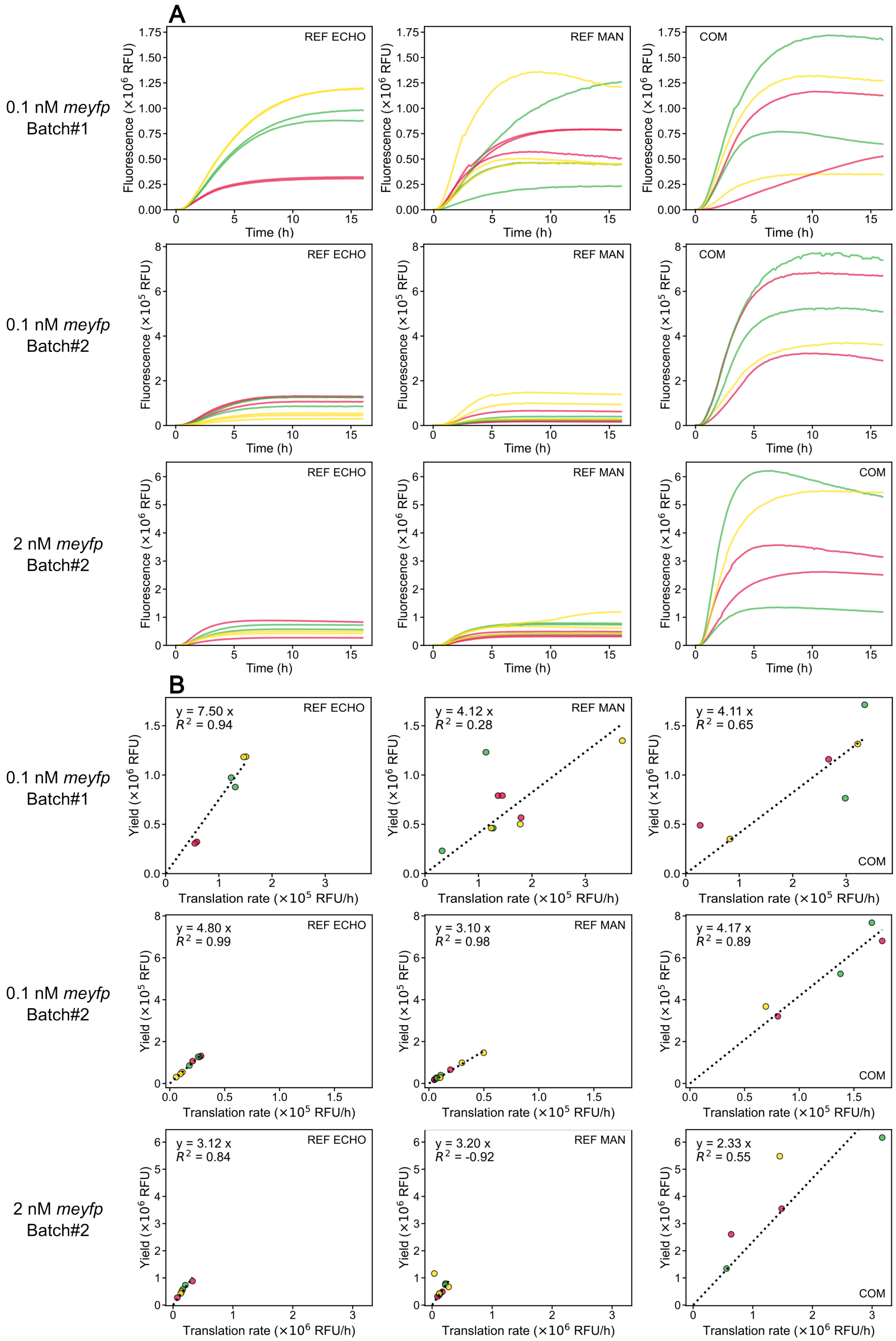


**Figure S1:** *legend on the next page*

**Figure S1: Kinetics of PURE reactions for different assembly methods and batches of components.** **(A)** Fluorescence kinetic traces of *meyfp* expression. **(B)** Scatter plots of apparent translation rate versus final protein yield as extracted from (A), for three PURE formulations: REF assembled with the Echo liquid handler (left column), REF assembled manually (middle column), and the commercial PURE*frex* 2.0 kit (right column). Rows correspond to (top) 0.1 nM DNA using Batch#1, (middle) 0.1 nM DNA using Batch#2, and (bottom) 2 nM DNA using Batch#2 of PURE components. In the kinetic plots (A), colors denote technical replicates of the same experimental condition; curves sharing the same color within a panel correspond to independent replicates of the same reaction condition. In the scatter plots, each dot represents one individual replicate, using the same color code as in the corresponding kinetic traces. Plotting kinetics, translation rate, and yield together enables direct comparison of performance and reproducibility across assembly methods, batches, and DNA concentrations. The linear fit and corresponding $R^2$ are shown. Overall, the commercial kit shows higher absolute yields, while Echo- and manually assembled REF formulations display comparable performance and reproducibility within each batch and DNA concentration.

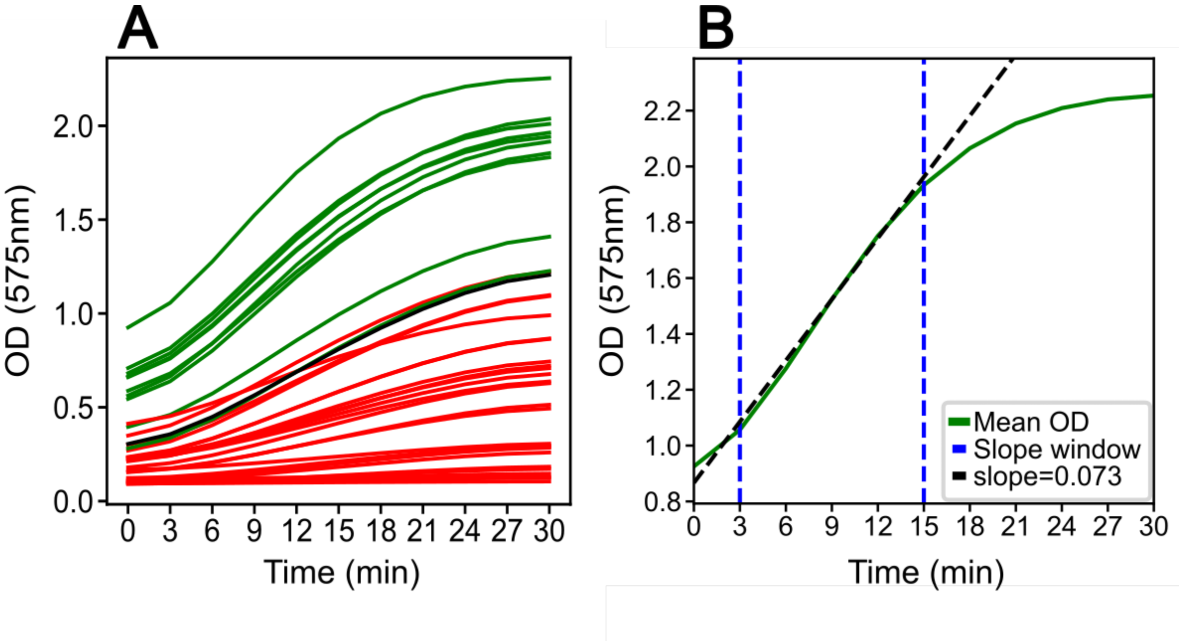


**Figure S2: Activity of expressed β-galactosidase as a second readout of PURE system performance. (A)** Mean absorbance kinetics at 575 nm for CPRG conversion in pre-ran PURE reactions (constant concentration of β-galactosidase) from round 2. Each curve represents the mean of 3 technical replicates. Green curves indicate conditions with higher activity than the reference (REF, black), while red curves indicate lower activity. **(B)** Extraction of the initial conversion rate through sliding window. The steepest slope window is highlighted (blue dashed lines) and the tangent line (black dashed) indicates the rate of CPRG conversion, which is proportional to the concentration of expressed β-galactosidase. This quantification allows direct comparison of the PURE system efficiency across different compositions.

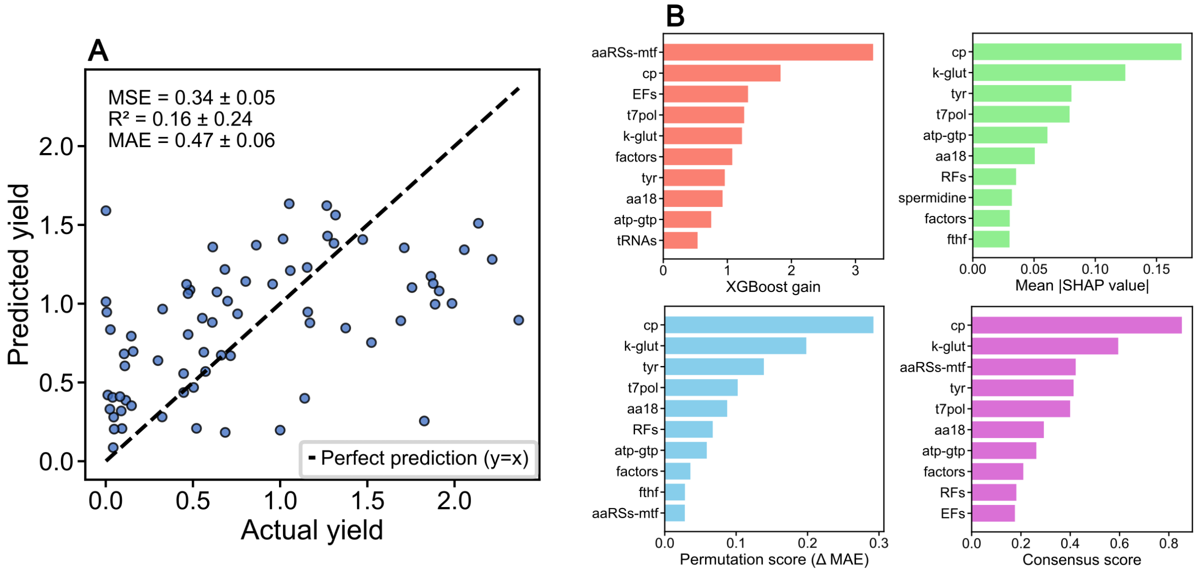


**Figure S3: XGBoost model performance and feature importance for yield prediction at 0.1 nM *meyfp* DNA. (A)** Predicted versus measured relative yields for the XGBoost model evaluated by nested cross-validation (CV = 5, five-fold cross-validation). Each point corresponds to one PURE composition. The dashed line indicates perfect prediction. Mean ± SD of MSE, $R^2$, and MAE across five outer folds are appended. **(B)** Feature importance metric comparison between XGBoost gain, SHAP values, permutation importance (ΔMAE), and a normalized consensus score. Bars show the top 10 components, identifying cp, k-glut, aaRS-mtf, t7pol, and tyrosine as major contributors to yield variation.

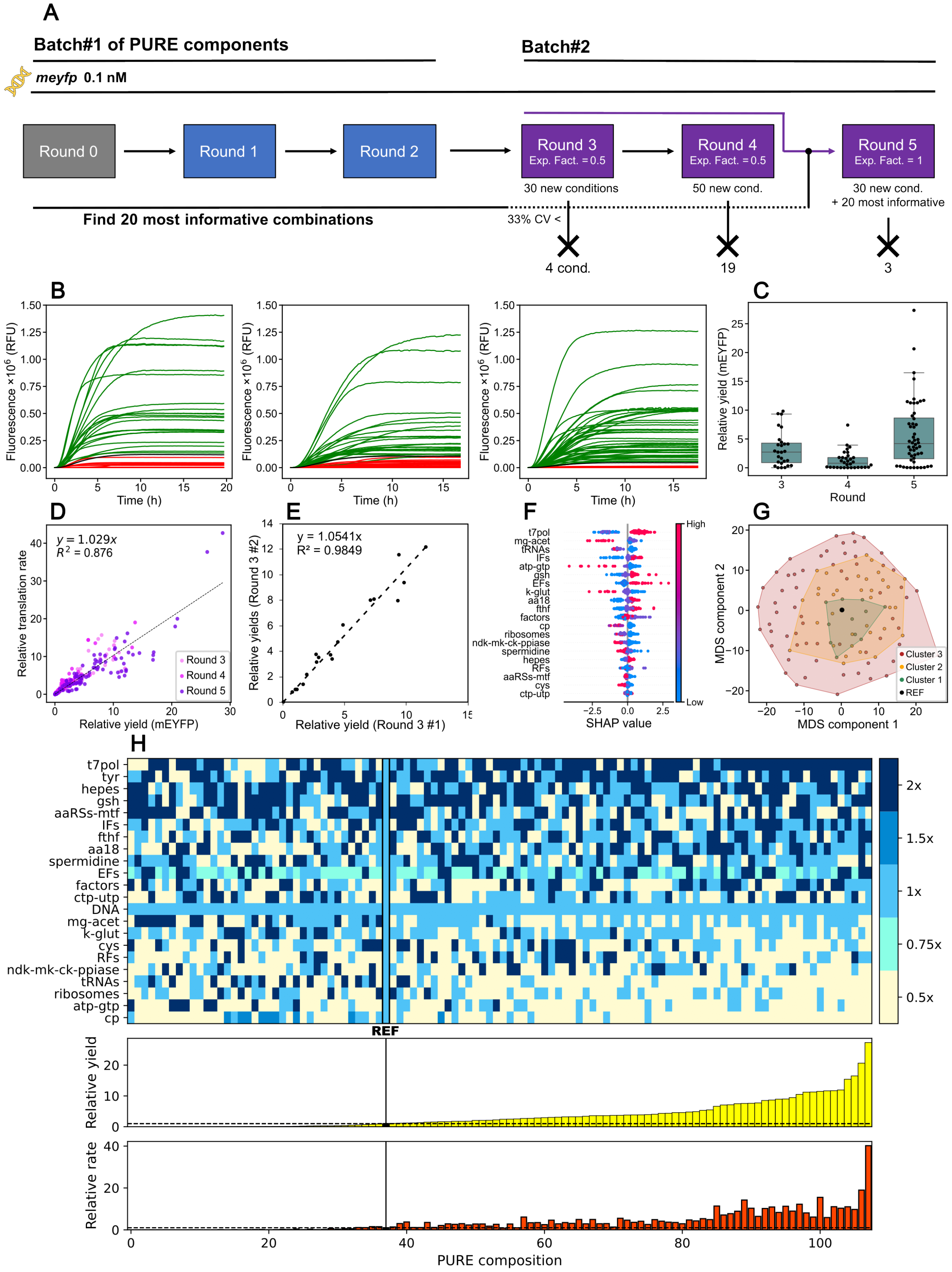


**Figure S4: Active learning consistently improves yield across component batches. (A)** Overview of conditions tested in rounds 3–5. Conditions with CV > 33% across technical replicates were discarded. Data from previous rounds were used to generate rounds 3 and 4. Round 5 was generated with data from rounds 3 and 4 and 20 most informative combinations from rounds 0 to 3. **(B)** Mean mEYFP fluorescence kinetics for all conditions tested across rounds 3–5. Each curve is the mean of two or three replicates. Black: REF composition (all components 1×). Green, yield > REF; red, yield < REF. **(C)** Distribution of relative expression yields for rounds 3–5. Box plots show median (center line), interquartile range (IQR, box), and 1.5× IQR (whiskers), with individual data points overlaid. **(D)** Correlation between relative translation rate and relative yield across rounds. **(E)** Reproducibility test of round 3. Relative yields from the initial round 3 experiment are plotted on the x-axis, and relative yields from a repeat experiment are plotted on the y-axis. Only relative yields of conditions that passed the CV% filtering in both experiments are shown here. The good correlation indicates the reproducibility of the same PURE reactions between experiments. The linear fit and corresponding $R^2$ are shown. **(F)** Shapley values (SHAP) beeswarm plot showing components most influencing yield prediction. **(G)** Multidimensional scaling (MDS) representation (Manhattan distances) of all conditions with k-means clustering (k = 3). High-yield conditions form a distinct, compact region, with nested hulls revealing a yield gradient. **(H)** Kymograph of all tested PURE conditions in rounds 3 to 5. High-yield conditions show elevated T7pol, tyr, and hepes, with lower cp, atp-gtp, tRNAs, and ribosomes, revealing distinct concentration patterns compared to lower-yielding compositions.

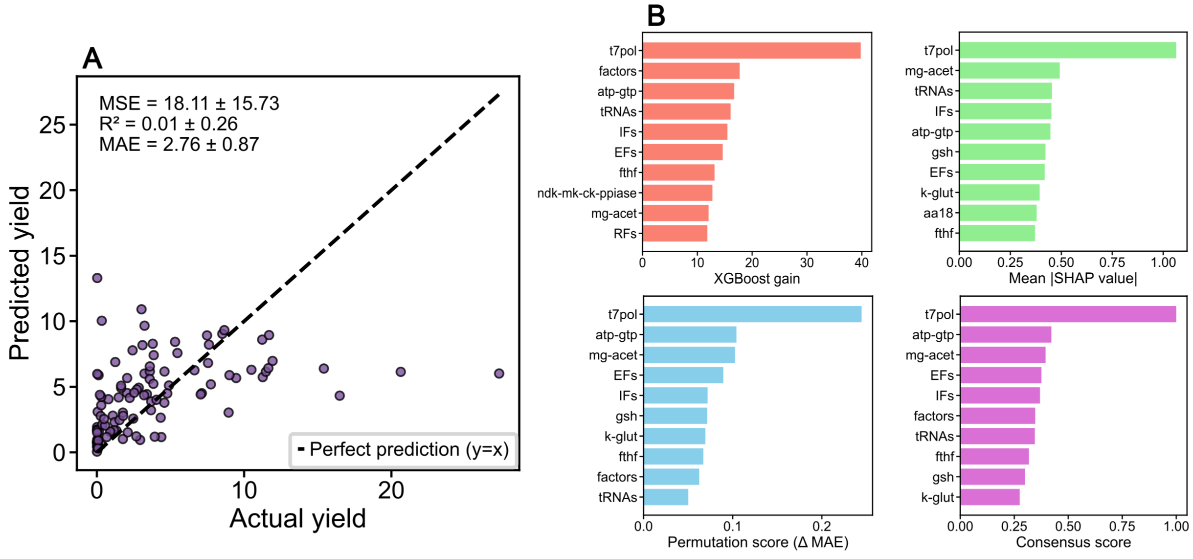


**Figure S5: XGBoost model performance and feature importance with the second batch of PURE components at 0.1 nM *meyfp* DNA. (A)** Predicted versus measured relative yields for the XGBoost model evaluated by nested cross-validation (CV = 5). Each point corresponds to one PURE composition. The dashed line indicates perfect prediction. Mean ± s.d. of MSE, $R^2$, and MAE across outer folds are appended. **(B)** Feature importance metric comparison between XGBoost gain, SHAP values, permutation importance (ΔMAE), and a normalized consensus score. Bars show the top 10 components, identifying t7pol as a major contributor to yield prediction.

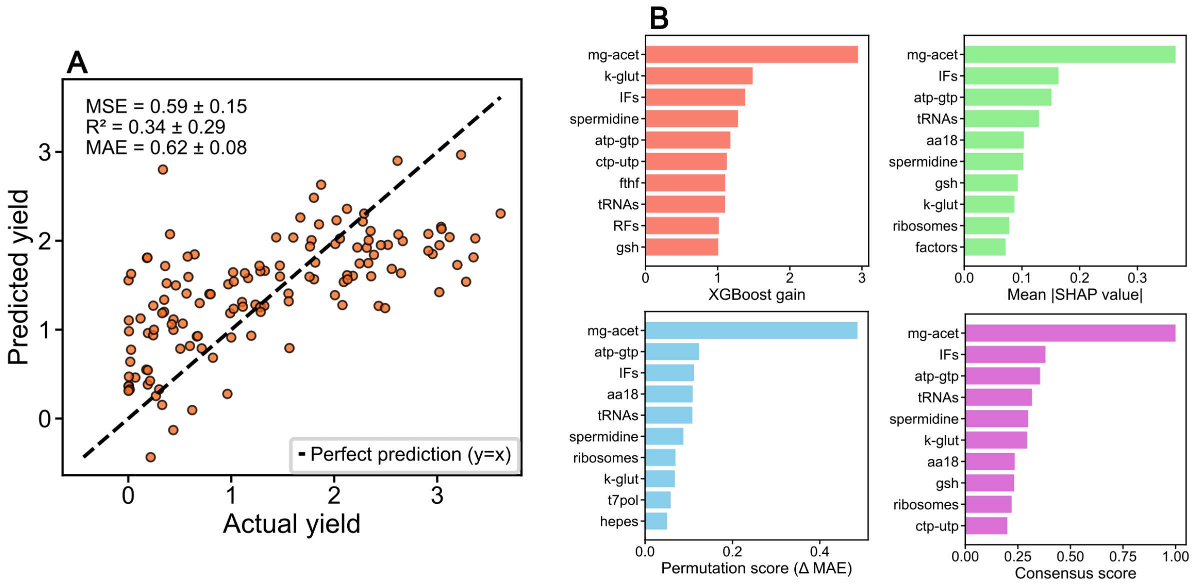


**Figure S6: XGBoost model performance and feature importance at 2 nM *meyfp* DNA. (A)** Predicted versus measured relative yields for the XGBoost model evaluated by nested cross-validation (CV = 5). Each point corresponds to one PURE composition. The dashed line indicates perfect prediction. Mean ± SD of MSE, $R^2$, and MAE across outer folds are appended. **(B)** Feature importance metric comparison between XGBoost gain, SHAP values, permutation importance (ΔMAE), and a normalized consensus score. Bars show the top 10 components, identifying mg-acet as a major contributor to yield prediction.

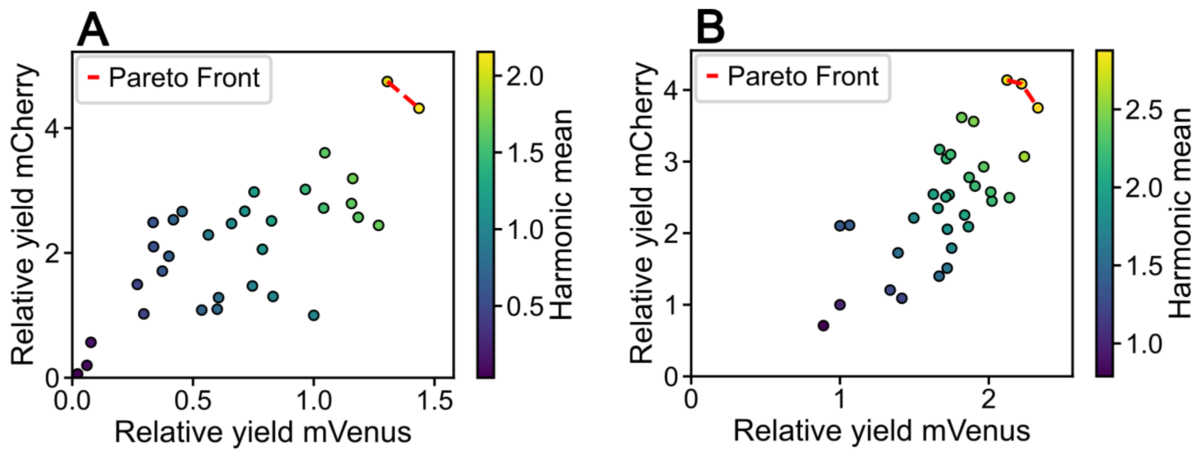


**Figure S7:** Scatter plots of relative mVenus versus mCherry yields for individual MSG1.1 optimization conditions at round 8 from 0.1 nM (**A**) or at 1 nM DNA (**B**). Colors indicate the harmonic mean of both reporter yields, and the dashed line marks the Pareto front for multi-objective optimization.

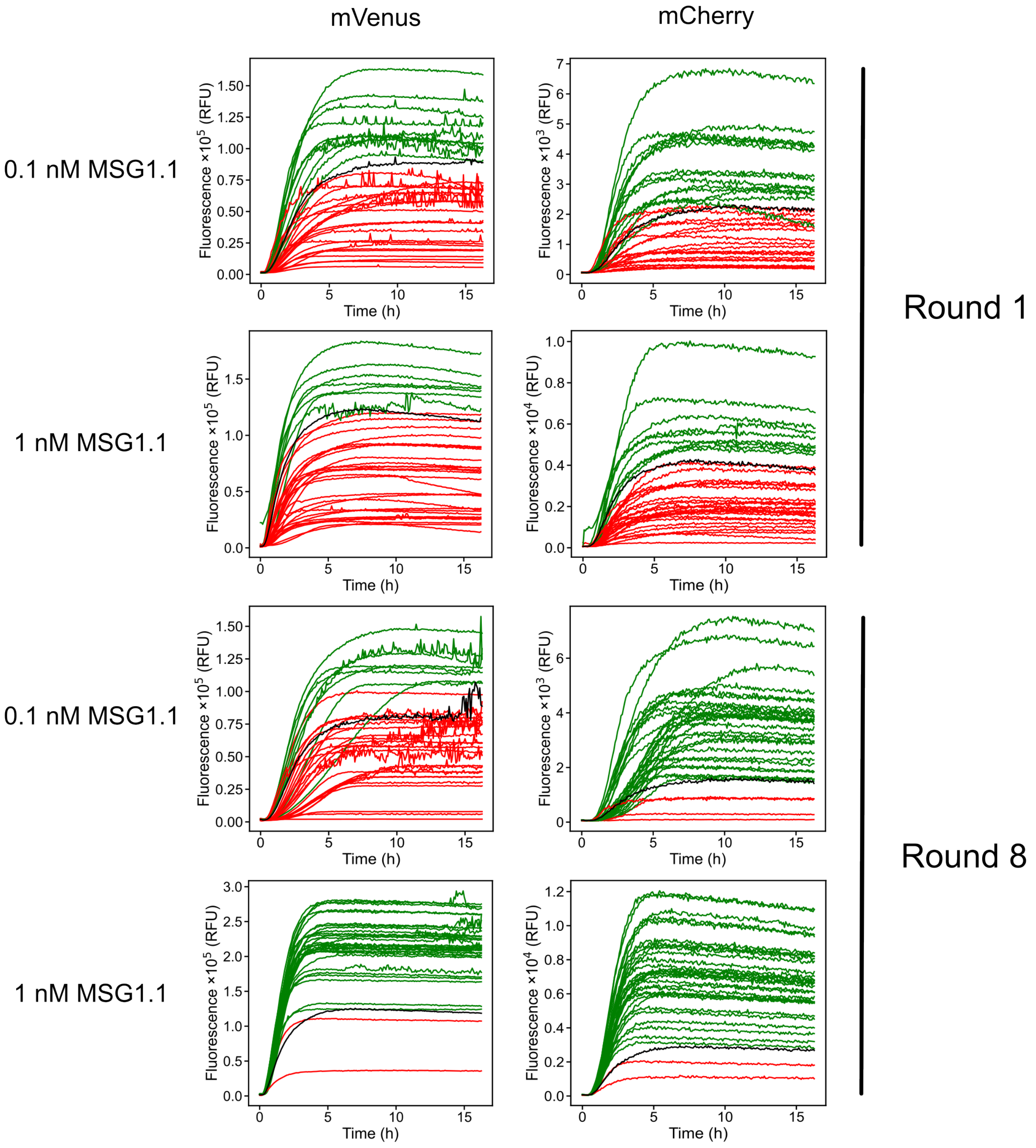


**Figure S8: PURE reactions kinetics for two different fluorescent reporters in rounds 1 and 8 of MSG1.1 expression optimization.** Mean mVenus and mCherry fluorescence kinetics for all conditions tested in rounds 1 and 8 at 0.1 nM and 1 nM of MSG1.1. Each curve is the mean of two replicates. Black: REF composition (all components 1×). Green, yield > REF; red, yield < REF.

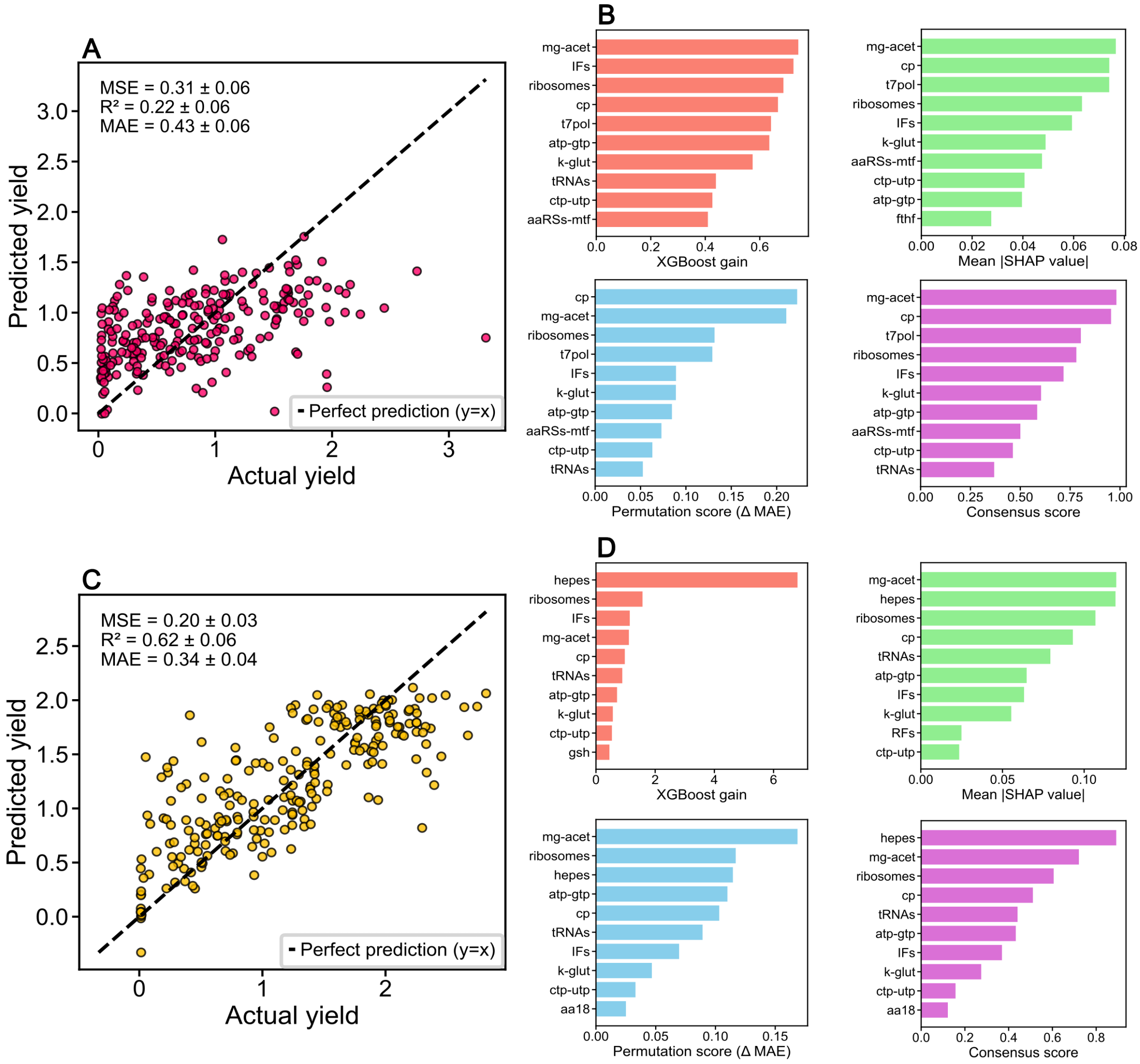


**Figure S9: XGBoost model performance and feature importance for MSG1.1 expression. (A)** Predicted versus experimentally measured harmonic mean for expression of MSG1.1 at 0.1 nM using nested 5-fold cross-validation. Each point corresponds to one PURE composition from the combined dataset, with the dashed line indicating perfect prediction. Performance metrics (MSE, $R^2$, MAE; mean ± SD across outer folds) are appended. **(B)** Feature importance analysis for the 0.1 nM DNA dataset, showing the top ten contributors ranked by XGBoost gain, mean absolute SHAP value, permutation importance (ΔMAE), and a normalized consensus score integrating all three metrics. **(C)** Predicted versus experimentally measured harmonic mean for expression of MSG1.1 at 1 nM DNA, assessed using the same nested cross-validation procedure as in **(A)**. The increased alignment of points along the identity line reflects improved model predictability at higher DNA concentration. **(D)** Feature importance analysis for the 1 nM DNA dataset, presented as in **(B)**. Across analysis methods, hepes, mg-acet, ribosomes, and cp consistently appear as key determinants of yield, with greater stability of rankings compared to the low-DNA condition.

| DNA | Component | Range | | | Stock |
|---|---|---|---|---|---|
| 0.1 nM | hepes | | | | 2× concentrated |
| | ribosomes | | | | - |
| | IFs | | | | 2× concentrated |
| | factors | | | | 2× concentrated |
| | k-glut | | | | - |
| | mg-acet | | | | 2× diluted |
| | Spermidine | | | | 2× diluted |
| | cys | | | | 2× diluted |
| | atp-gtp | | | | - |
| | EFs | | | | 2× diluted |
| | RFs | | | | 2× diluted |
| | t7pol | | | | 2× diluted |
| 1 nM | tRNAs | | | | - |
| | ribosomes | | | | - |
| | IFs | | | | 2× concentrated |
| | k-glut | | | | - |
| | mg-acet | | | | 2× diluted |
| | spermidine | | | | 2× diluted |
| | aa18 | | | | - |
| | atp-gtp | | | | - |
| | gsh | | | | - |
| | cp | | | | - |

4×
3×
2×
1×
0.5×
0.25×

**Table S1:** Concentrations of components utilized in rounds 6-8 for MSG1.1 expression optimization. 1× indicates a concentration identical to that in the regular PURE*frex* 2.0 kit.

| Plasmid name | Characteristics |
|---|---|
| G76 | pUC57 parental plasmid<br>ori; AmpR; ColE1<br>*meyfp* transcriptional unit: pT7- *meyfp* -LL-spinach T7t<br>(codon optimized) |
| pT003 | pUC57 parental plasmid<br>ori-*lacZ*; AmpR; ColE1<br>*lacZ* transcriptional unit: pT7-g10L-RBS-lacZ-T7t |
| MSG1.1 | CEN6/ARS4; pT7-p2-tVSV-tVSV; pT7-p3-tT7; pHHF2-crtE-tADH1; pSP6-eYFP-LL-spinach-tT7; pTDH3-crtI-tTDH ARS417; pT7-minD-tT7; pT7-minE-tT7; pCCW12-mRuby2-tENO1; pT7-ftsZ: mVenus-ftsA-tT7; pTEF-hphNT1-tTEF; oriR-oriL; cat; redF; oriV; ori2; repE; incC; sopA; sopB; sopC; pURA3-URA3-tURA3; pT7-mCherry-tT7; pPGK1-crtYB-tPGK1; pT7-plsB-tT7 pT7-plsC-tT7; pT7-cdsA-tT7; pT7-pssA-tT7; pT7-psd-tT7; pSP6-pgsA-tT7; pSP6-pgpA-tT7 |

**Table S2:** List of plasmids used in this study. pT7 = T7 promoter, tT7 = T7 terminator, tVSV = VSV terminator, pSP6 = SP6 promoter, LL = long linker, RBS = ribosome binding site.

| Primer | DNA sequence (5'-3') | Purpose |
|---|---|---|
| M13F | CCCAGTCACGACGTTGTAAAACG | Amplify *meyfp* expression cassette from G76 |
| M13R | AGCGGATAACAATTTCACACAGG | |
| ChDT-16 | AAAGTAAGCCCCCACCCTCACATG | Amplify *lacZ* expression cassette from pT003 |
| ChDT-17 | AAAGTAGGGTACAGCGACAACATACAC | |
| IVIV_BB_fw | CCAGGTTTTCACCGTAACAC | Amplify pCC1BAC marker from MSG1.1 |
| pCC1BAC_BB_dg_rv | gaatggcggaacgtatgtgg | |
| pCCW2_dg_FW | CCGTTCAAGTATATAAAGTCGGC | Amplify mRuby2 marker from MSG1.1 |
| ENO1ConRV | ACATGGGTGACCAAAAGAGC | |
| CEN6/ARS4_FW | atcacgtgctataaaaataattataatttaaat | Amplify CEN6/ARS4 marker from MSG1.1 |
| FK161 | GATCGCTTGCCTGTAACT | |

**Table S3:** List of primers used in this study.

| Hyperparameter | Description | Values explored | Values selected | | | | |
|---|---|---|---|---|---|---|---|
| | | | A | B | C | D | E |
| *learning_rate* | Step size shrinkage used to prevent overfitting | 0.01, 0.03, 0.1, 0.3 | 0.3 | 0.03 | 0.3 | 0.03 | 0.01 |
| *colsample_bytree* | Fraction of features randomly sampled for each tree | 0.6, 0.8, 0.9, 1.0 | 0.8 | 0.8 | 0.9 | 0.9 | 0.6 |
| *subsample* | Fraction of training samples randomly sampled for each boosting iteration | 0.6, 0.8, 0.9, 1.0 | 1 | 0.8 | 0.9 | 0.8 | 0.9 |
| *max_depth* | Maximum depth of individual trees | 2, 3, 4, 6, 8 | 4 | 8 | 4 | 8 | 6 |
| *n_estimators* | Number of boosting trees | 10, 20, 40, 60, 80, 100, 300, 500 | 40 | 500 | 80 | 100 | 500 |
| *reg_lambda* | L2 regularization term on weights | 1, 1.5, 2 | 2 | 1.5 | 1.5 | 1 | 2 |
| *gamma* | Minimum loss reduction required to make a further partition on a leaf node | 0, 0.1, 0.4, 0.6 | 0.1 | 0.6 | 0.1 | 0 | 0.1 |
| *min_child_weight* | Minimum sum of instance weights needed in a child node | 1, 2, 4 | 2 | 1 | 2 | 4 | 4 |

**Table S4:** List of hyperparameters used in METIS/ML. The selected values correspond to those used to train the final XGBoost model, obtained as the most frequently selected hyperparameters across the outer folds of a nested 5-fold cross-validation with randomized hyperparameter search. Experimental conditions: A) 0.1 nM *meyfp* Batch#1, B) 0.1 nM *meyfp* Batch#2, C) 2 nM *meyfp* Batch#2, D) 0.1 nM MSG1.1, and E) 1 nM MSG1.1.
